# Exosomes regulate Neurogenesis and Circuit Assembly in a Model of Rett Syndrome

**DOI:** 10.1101/168955

**Authors:** Pranav Sharma, Pinar Mesci, Cassiano Carromeu, Daniel McClatchy, Lucio Schiapparelli, John R Yates, Alysson R. Muotri, Hollis T Cline

## Abstract

Exosomes are thought to be secreted by all cells in the body and to be involved in intercellular communication. Here, we tested whether neural exosomes regulate the development of neural circuits and whether exosome-mediated signaling may be aberrant in the neurodevelopmental disorder Rett Syndrome (RTT). Quantitative proteomic analysis comparing exosomes from human induced pluripotent stem cells (hiPSC) - derived RTT patient or control neural cultures indicates that control exosomes contain signaling components capable of influencing neuronal development and function, which are lacking in RTT exosomes. Moreover, treatment with control exosomes rescues neuron number, apoptosis, synaptic puncta and synchronized firing phenotypes of MeCP2 knockdown in human primary neurons, indicating that exosomes have the capacity to influence neural development and may be a promising avenue to treat neurodevelopmental disorders like Rett Syndrome.

**Highlights:** Exosome proteomics distinguish cargo in RTT vs control hiPSC-derived neural cultures Control but not RTT exosomes increase neurogenesis in human neural cultures hiPSC-derived neural exosomes reverse pathological phenotypes in RTT neural cultures RTT exosomes do not impair neural development

## Introduction

The nervous system is composed of one of the most complex dynamic multicellular tissue architectures in the body and requires short- and long-range intercellular communication for its development and function. Exosomes are small vesicles secreted by all cells in the brain, including neurons, and have been hypothesized to play a critical role in cell-cell communication (Budnik et al., 2016; Sharma et al., 2013). Exosomes carry protein, lipid and RNA cargo and have the capacity to signal to recipient cells to modify biological functions. Exosomes can signal over short range within brain tissue (Zappulli et al., 2016), and can signal widely throughout the brain through the cerebrospinal fluid (Chiasserini et al., 2014; Coleman and Hill, 2015). The ability of exosomes to cross the blood-brain barrier extends the putative range for their bioactive influence to the entire organism (Alvarez-Erviti et al., 2011; Graner et al., 2009). Strong evidence indicates that exosomes impart biological activity to neurons (Rajendran et al., 2014; Zappulli et al., 2016). For instance, in the *Drosophila* larval neuromuscular junction, exosome-mediated protein transport is required for coordinated development of pre- and postsynaptic components of the neuromuscular junction (Korkut et al., 2009; Korkut et al., 2013). Exosomes secreted by oligodendrocytes affect firing rate, signaling pathways and gene expression in cultured primary neurons (Frohlich et al., 2014; Fruhbeis et al., 2013). While there is evidence of biological roles of exosomes secreted by neurons and other cell types in brain, their function in the development of human neural circuits is largely unexplored.

The detection of proteins associated with neurodegenerative diseases in exosomes led to increased interest in their role in disease (Budnik et al., 2016; Howitt and Hill, 2016; Sarko and McKinney, 2017). Exosomes have been implicated in the progression of neurodegenerative proteinopathies, based on the transcellular transfer of proteins like misfolded prion protein in Creutzfeldt-Jakob disease (CJD), amyloid β peptide and tau/phosphorylated tau in Alzheimer’s disease (AD), SOD1/mutant SOD1 and TDP-43 in amyotrophic lateral sclerosis (ALS), and α-synuclein in Parkinson’s disease (PD) (Budnik et al., 2016; Howitt and Hill, 2016; Loov et al., 2016; Sarko and McKinney, 2017). Similarly, exosomes secreted by microglia and astrocytes transfer pro-inflammatory signals in multiple sclerosis (MS) (Carandini et al., 2015). Exosomes also play a profound role in cancers, including glioblastomas (Budnik et al., 2016; Gourlay et al., 2017), however their proposed function in transferring pathogenic proteins in neurodegenerative diseases contrasts sharply with their role in cancer. Exosomes released by cancer cells signal to diverse target tissues to facilitate development and progression of tumors (Gourlay et al., 2017; Hoshino et al., 2015; Zhang et al., 2015).

Rett syndrome (RTT) is a progressive neurodevelopmental disorder primarily caused by loss of function of the X-linked *Methyl*-*CpG-binding protein 2* (*MeCP2*) gene (Amir et al., 1999; Feldman et al., 2016; Leonard et al., 2017; Zoghbi, 2016). RTT patients display a distinct clinical signature where infants exhibit normal early growth and development up to 6-18 months, followed by regression of the acquired abilities. Affected individuals display a range of clinical symptoms depending on the severity of disease that includes acquired microcephaly, loss of purposeful hand use, stereotyped movements, including hand wringing, gait apraxia and ataxia, tremor, seizures, apnea, hyperventilation, severe intellectual disability and features of autism spectrum disorder (ASD). While the primary role of MeCP2 is transcriptional repression through DNA methylation, it is also involved in chromatin remodeling, gene activation, regulation of alternative splicing, and microRNA (miRNA) processing (Ausio et al., 2014; Zoghbi, 2016). Although RTT CNS phenotypes include deficits in neuronal circuit development where intercellular communication could be important (Zoghbi, 2016), thus far the role of exosomes has not been explored. Specifically, it remains unclear whether exosomes actively facilitate development of disease pathology in RTT, or conversely, whether exosomes from healthy cells might provide corrective measures to ameliorate abnormalities resulting from disease.

To investigate the role of exosomes in neural circuit development, we developed a reductionist experimental paradigm in which we added purified exosomes isolated from human induced pluripotent stem cells (hiPSCs)-derived neurons onto human primary neural cultures to assay their capacity to influence neuron and circuit development. We found that treatment with exosomes increased neurogenesis by promoting cell proliferation and neuronal differentiation, suggesting that exosomes carry signaling components that influence cell fate in developing neural circuits. We have previously shown that neurons derived from RTT-hiPSC have fewer synapses, reduced spine density, smaller soma size, altered calcium signaling and electrophysiological defects compared to control neurons (Marchetto et al., 2010). We took advantage of this human disease model to investigate the role of exosomes in developing neural circuits, using RTT patient hiPSC-derived neural cultures (RTT neural cultures) and a CRISPR/Cas9 isogenic rescue of the *MECP2* mutation in the patient cell line hiPSC-derived neural cultures (control neural cultures) (de Souza et al., 2017; Zhang et al., 2016). We purified exosomes released from RTT neural cultures and compared them with exosomes from isogenic control neural cultures by quantitative proteomic analysis. RTT exosomes differed significantly from control exosomes in their protein signaling content and were predicted to elicit phenotypic outcomes, such as deficient synaptic connectivity, consistent with RTT. Therefore, we tested the hypothesis that treatment with control exosomes could rescue RTT-associated phenotypes in human primary neural cultures with MeCP2 knock down. While knocking down MeCP2 in human primary neurons with short hairpin RNA (shRNA) increased apoptosis, decreased the number of synapses and decreased synchronous firing of neurons, treatment with control exosomes rescued these phenotypes. Thus, we show for the first time that exosomes play a profound role in neuronal circuit development and can be used to reverse deficits in neurological disease models.

## Results

### Exosome purification from human iPSC-derived neural cultures

To test the hypothesis that exosomes play a physiological role in neuronal development, we established neural cultures derived from hiPSCs as a source for exosomes, because of the demonstrated value of iPSCs in modeling human diseases (Zhang et al., 2016, Marchetto et al., 2010, de Souza et al., 2017), and the capacity to harvest large volumes of exosomes. We cultured hiPSC-derived neurons in serum-free growth and differentiation media with verified components to avoid potential contamination from external exosomes. Exosomes were purified using a differential centrifugation protocol (Thery et al., 2006), slightly modified for our cultures (Figure 1A). Electron microscopic analysis demonstrated that purified exosomes secreted by neural cultures of hiPSCs had typical disc shaped morphology (Figure 1B) and ranged in size from 30-200 nm; ∼60% of exosomes were between 40-80 nm and >80% were <120 nm (data not shown). Western blot analysis showed that the purified exosomes contain the typical exosome markers, Alix and Flotilin, and that RTT exosomes had less Alix than control exosomes, compared to the loading control, β-actin (Figure 1C), suggesting that RTT and control exosomes may differ in their protein cargo.

**Figure 1:**
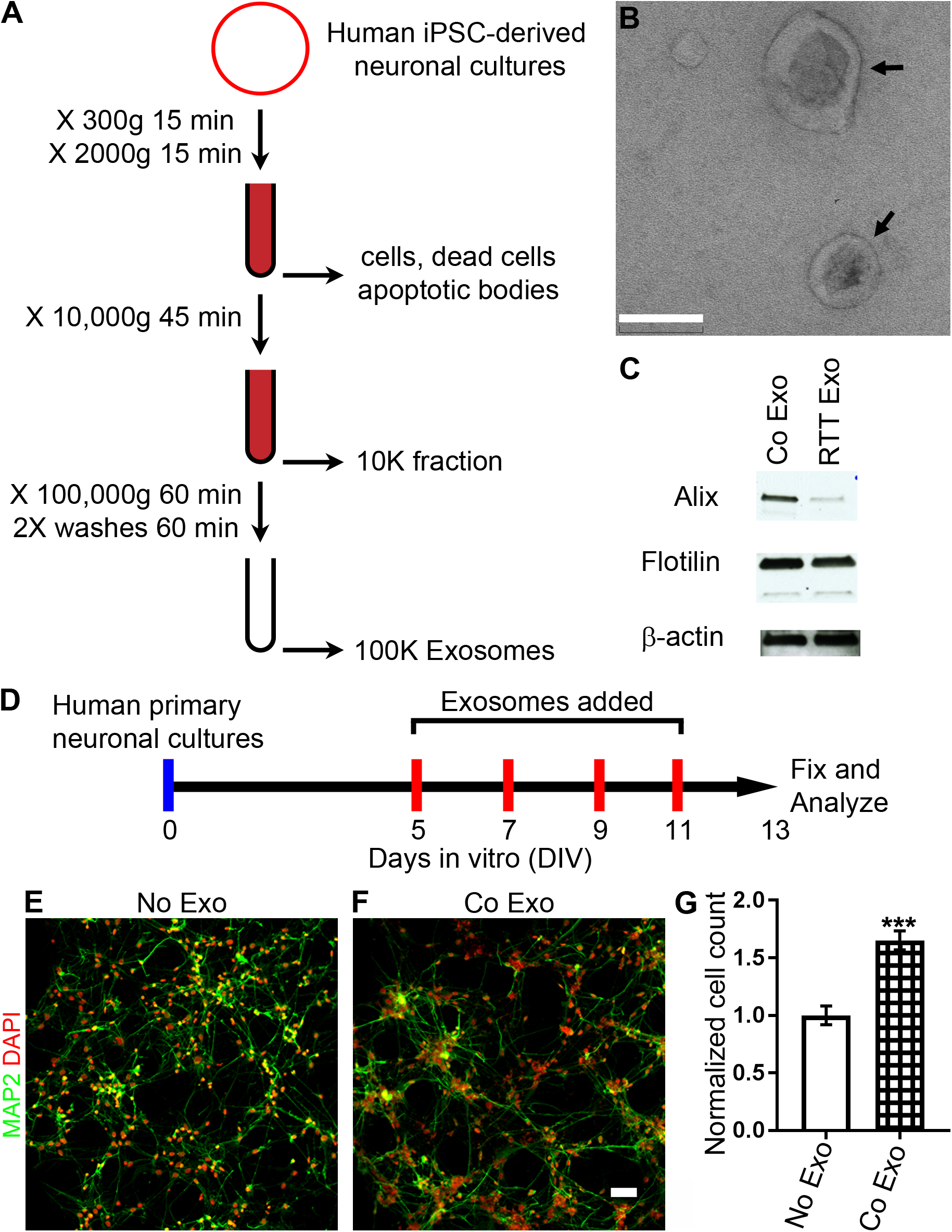
Exosomes increase cell number in developing neural cultures. A. Protocol for purification of exosomes using ultracentrifugation. B. Electron micrograph of purified exosomes, arrows. C. Western blot analysis showing exosomal markers, Alix and Flotilin, in exosomes purified from control and RTT human iPSC-derived neural cultures. D. Protocol for treatment of human primary neural cultures with purified control exosomes (Co Exo) or an equal volume of media (No Exo) added on DIV 5, 7, 9 and 11. Cultures were fixed on DIV 13, labeled and analyzed. E, F. Images of human neural cultures treated with media (No Exo, E) or control exosomes (Co Exo, F), labeled with MAP2 antibodies (green) or DAPI (red). G. Graph of the total cell number in cultures treated with control exosomes compared to media. Exosome treatment increased total cell number 1.65X ± 0.08 (*p*=0.001, n=4 wells per group, 2 way ANOVA). Scale bar = 0.1 μm (B), 20 μm (F).

### Exosomes increase cell number in developing neural cultures

To investigate the effect of exosomes on developing neurons we established a protocol for treating neural cultures with exosomes. Donor neural cultures derived from hiPSCs were differentiated for 6 weeks from neural progenitor cells (NPCs) to ensure maturation of neurons (Marchetto et al., 2010). Six weeks after plating, media conditioned for 2 days was harvested and used for purification of exosomes as described in Figure 1A. Purified exosomes were resuspended in fresh growth media and applied to recipient neuronal cultures. An equivalent volume of fresh growth media without exosomes applied to control cultures. We exposed hiPSC-derived neural cultures as well as human primary neural cultures to exosomes assess the effects of exosome treatments. The human primary neural cultures provide the benefit of a shorter span of maturation compared to iPSC-derived neural cultures. To assess the effect of exosome treatment on cell number, we exposed neural cultures to exosomes, starting on 5 days *in vitro* (DIV) and adding exosomes every two days until 11 DIV (Figure 1D). We fixed the recipient primary human neural cultures on day 13 and analyzed for total cell number. The human primary neuron cultures show widespread MAP2 expressing neurons with typical neuronal morphologies by 13 DIV, indicating healthy differentiated cultures (Figure 1E-F). Cell number, determined by DAPI labeling, was significantly greater in exosome-treated cultures compared to control (1.65X ± 0.08, *p*=0.001, Figure 1E-G).

### Exosomes contain signaling proteins that are altered in the absence of MeCP2

Since exosomes increased the number of cells in developing cultures, and because changing cell number is an important hallmark of neural circuit maturation, we investigated exosome function in developing neural cultures and in the context of the neurodevelopmental disorder, Rett syndrome. First, we conducted a quantitative mass spectrometry proteomic study of exosomes using TMT isobaric tags, as an unbiased approach to analyze exosome proteins. Exosomes isolated from hiPSC-derived neural cultures from a male RTT patient deficient in *MECP2* expression (RTT exosomes) were compared to exosomes isolated from isogenic hiPSC-derived neural cultures that were corrected for the *MECP2* mutation using CRISPR/Cas9 (control exosomes) (Zhang et al., 2016). We detected a total of 2572 proteins from RTT and control exosomes in two independent MS experiments (Table S2, tab 1), of which 739 were annotated for function in neurons. Analysis of ‘cellular pathways’ and ‘downstream effects on biological functions in the nervous system’ using the manually curated Ingenuity database (Calvano et al., 2005) indicated that exosomes secreted by hiPSC derived neural cultures are enriched in proteins that impact neuronal development and function. Specifically, they contain signaling components for protein translation (EIF2, eIF4), axonal guidance, integrin signaling, ephrin signaling, and cytoskeletal regulation (Rho family, Actin cytoskeleton, Rho GDI), among the most significant categories (Figure S2C, Table S4, tab 2), and had capacity to influence downstream functions like neuritogenesis, development, morphogenesis and proliferation of neurons, and synaptic development and function (Figure S2D, Table S5, tab 2).

**Figure 2:**
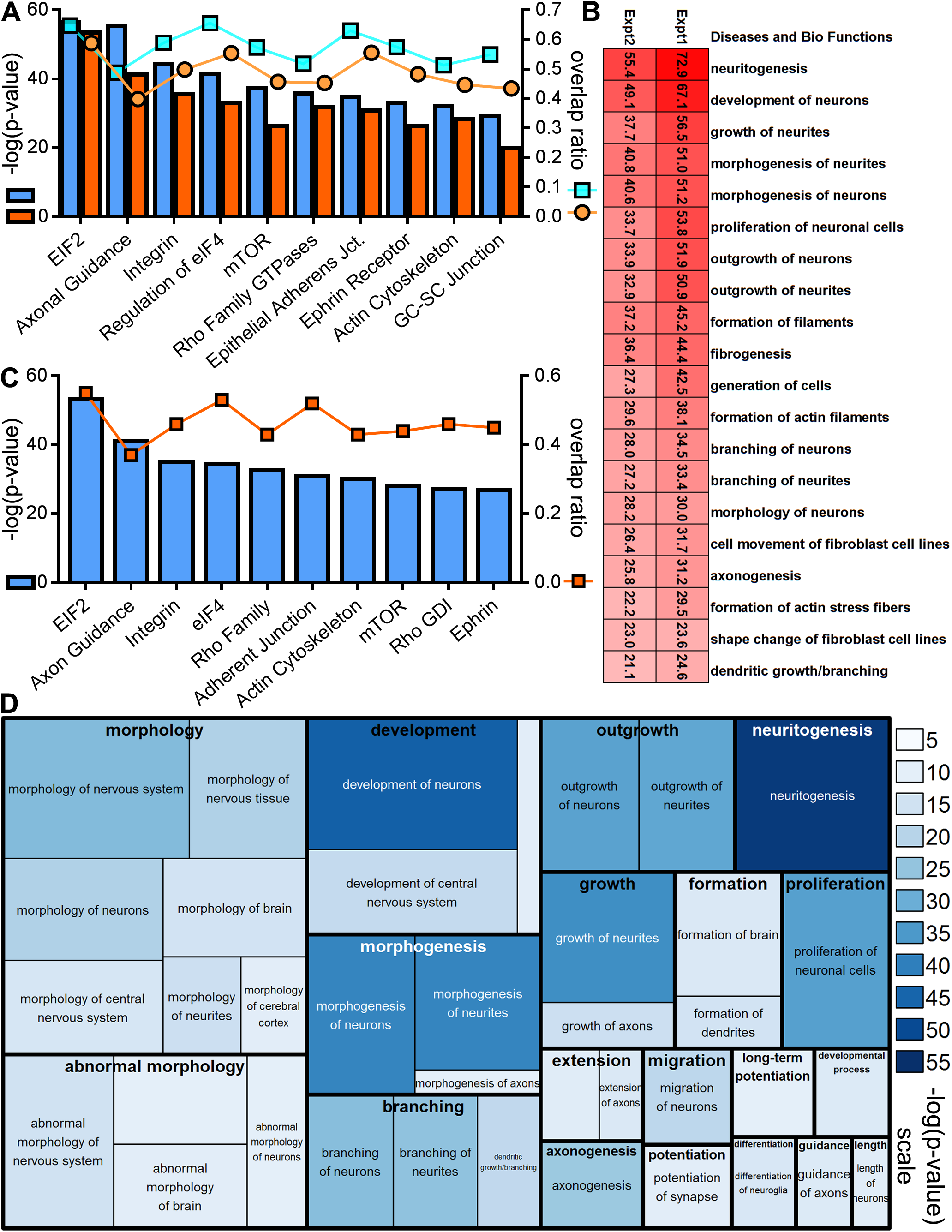
Exosomes contain complex signaling components that are altered with MeCP2 mutation. A. Pie charts showing the cellular distribution of proteins identified in quantitative proteomic analysis of exosomes isolated from control and RTT patient iPSC-derived neural cultures based on annotation using Ingenuity. The dataset of 237 proteins with >1.5 fold difference between control and RTT exosomes was enriched in plasma membrane proteins compared to the dataset of all 2572 exosome proteins. B. PANTHER Overrepresentation Test of the dataset of 237 proteins with >1.5 fold change using the GO ‘biological processes’ complete annotation dataset. The 237 candidates were classified into annotated GO biological process categories and compared with the normal human database to determine whether they are overrepresented or underrepresented for a given GO biological process. Fold enrichment (X-axis) is the ratio of proteins classified in each GO category from the experimental dataset over the number of proteins expected in the GO category predicted from the reference normal human dataset. Positive values of fold change indicate overrepresentation of the GO category in the experimental dataset. The top 10 GO biological processes by p-values were plotted as a heatmap, where color intensity depicts -log(p-value) from 15.98 (bright red) to 10.38 (light red). ‘Neurogenesis’ and ‘nervous system development’ are among the most significantly enriched GO categories. C. Top 10 canonical pathways from Ingenuity analysis using the dataset of 237 proteins. Blue bars, associated with the left Y axis show significance values plotted as -log(p-value) and the square orange markers, associated with the right Y axis show the overlap ratios of the number of proteins from our dataset relative to the total number of proteins annotated to each canonical pathway. D. Treemap heatmap showing Ingenuity analysis of the relative strength of downstream effects for biological functions and diseases in the nervous system using the dataset of 237 proteins. The predicted biological functions are plotted hierarchically as a treemap, where the box size represents the number of proteins from dataset annotated for given function, ranging from 2 to 44 proteins, and the heatmap with the color look up table indicates significance values plotted as -log(p-values) at right.

Out of the 2572 proteins present in both RTT and control exosomes, 237 had an average fold difference of 1.5 or more between RTT and control samples (Table S2, tab 2). When we compared these 237 differentially packaged exosome proteins with the entire dataset of exosome proteins, we found that they were enriched with proteins annotated to be in the plasma membrane (38% compared to 19% for all proteins) whereas cytoplasmic proteins were relatively underrepresented (31% compared to 51%; Figure 2A). We analyzed this dataset of 237 proteins using the Ingenuity database and the PANTHER Overrepresentation Test (Mi et al., 2013) and GO enrichment using FunCoup (Alexeyenko and Sonnhammer, 2009). The PANTHER Overrepresentation Test using the GO ‘biological processes’ complete annotation dataset indicated that ‘nervous system development’ (+2.88 fold enrichment, *p* = 4.96 × 10^−13^) and ‘neurogenesis’ (+3.26 fold enrichment, *p* = 4.40 × 10^−11^) were highly overrepresented compared to the normal human database (Figure 2B, Table S3). Ingenuity analysis of the representation of components with a role in different cellular functions indicated that ‘axon guidance’ (p = 3.62 × 10^−8^), ‘ephrin signaling’ (Ephrin B: *p* = 1.62 × 10^−4^, Ephrin receptor: *p* = 7.24 × 10^−4^) and ‘actin dynamics’ (actin nucleation: *p* = 3.89 × 10^−04^, RhoGDI: *p* = 7.24 × 10^−4^) are among the most significant (Figure 2C, Table S4, tab 1). While many pathways that were predicted based on the dataset of differentially packaged exosome proteins are similar to those predicted based on the dataset of all proteins, key interesting differences are in ‘SUMOylation’ (*p* = 1.17 × 10^−7^), ‘glioblastoma multiforme (GBM) signaling’ (*p* = 1.17 × 10^−7^) and ‘high mobility group box 1 (HMGB1) signaling’ (*p* = 1.17 × 10^−7^). SUMOylation is involved in neuronal differentiation as well as development and maintenance of synapses (Henley et al., 2014). MeCP2 itself is SUMOylated and SUMOylation is critical for the transcriptional repression function of MeCP2 (Cheng et al., 2014; Tai et al., 2016). HMGB1 is widely expressed in the brain during development and adulthood and plays a role in neurite outgrowth, cell migration and neuroinflammation (Fang et al., 2012). Furthermore, HMGB1 binds MeCP2 in biochemical assays (Dintilhac and Bernues, 2002). Exosomes released from GBM are thought to modify cells or extracellular substrates in the microenvironment to aid tumor progression (Gourlay et al., 2017). GBM signaling components identified here in neural exosomes include regulators of cytoskeletal machinery which could have similar roles in altering interactions between developing neural cells and the microenvironment in Rett Syndrome (Gourlay et al., 2017).

The Ingenuity Pathway Analysis (IPA) curated ‘downstream effects on biological functions in the nervous system’ predicted roles in ‘neuronal development’ (*p* = 1.89 × 10^−13^), ‘neuritogenesis’ (*p* = 9.47 × 10^−12^), ‘axonal guidance’ (*p* = 1.35 × 10^−11^), ‘proliferation of neurons’ (*p* = 1.25 × 10^−9^) and ‘synaptic development’ (*p* = 3.08 × 10^−7^) among the most significant categories (Figure 2 D). However, the ‘abnormal morphology of nervous system’ category stands out with respect to known phenotypes of *MECP2* mutation and RTT (Figure 2 D, Table S5, tab 1).

To summarize, quantitative proteomic and bioinformatic analyses indicate that exosomes contain a range of signaling components that are capable of influencing neural development, as well as neuron and circuit function. The protein signaling codes affecting multiple aspects of neural development are altered in exosomes released from hiPSC derived neural cultures lacking MeCP2 protein in broadly classified categories of proliferation, neurogenesis and synapse development and function. We test the predicted functions of exosomes in neural development in the experiments below.

### Control exosomes increase proliferation and survival in developing neural cultures while RTT exosomes have no effect

The proteomic analysis above indicates that RTT and control exosomes may have different effects on cell proliferation and neural development. To test these possibilities we labeled dividing cells with 5-ethynyl-2′-deoxyuridine (EdU), a thymidine analog that is incorporated into DNA in cells undergoing active DNA synthesis during S-phase of cell cycle, to study cell proliferation. We incubated primary human neural cultures with 10 μM of EdU for 4 hours and analyzed EdU immunolabeling on DIV 5,7,9, and 11. Some of the EdU labeled cells displayed typical NPC morphology and expressed Nestin, an intermediate filament protein that is a marker of NPCs (Figure S3A). We observed EdU labeled cells at all time points, demonstrating that the cultures maintain active proliferation until DIV 11, however proliferation was significantly reduced on DIV 11 (Figure S3B). Based on these results, we tested the effect of exosome treatments on DIV 5 and 7 on cell proliferation and differentiation by treating human neural cultures with media alone (no exosomes), control exosomes or RTT exosomes on DIV 5 and 7 (Figure 3A). On day 7, a 2-hour pulse of 10 μM EdU was provided prior to addition of exosomes. The cultures were maintained until DIV 9, then assayed to quantify EdU-labeled NPC progeny. Control exosome treatment expanded the progeny of EdU labeled progenitors by a factor of 1.35 ± 0.09 (*p*=0.006) compared to no exosome treatment (Figure 3 B-E), whereas the effect of treatment with RTT exosomes was not statistically different from the addition of media alone. These results show that control exosomes contain signals that promote NPC proliferation and survival of EdU-labeled progeny. These proliferation and survival signals are lacking in RTT exosomes from hiPSC-derived neural cultures lacking MeCP2 protein. These results are consistent with our proteomic data analysis where proteins that were significantly altered between control and RTT exosomes were predicted to play a role in neuronal proliferation (Figure 2D).

**Figure 3.**
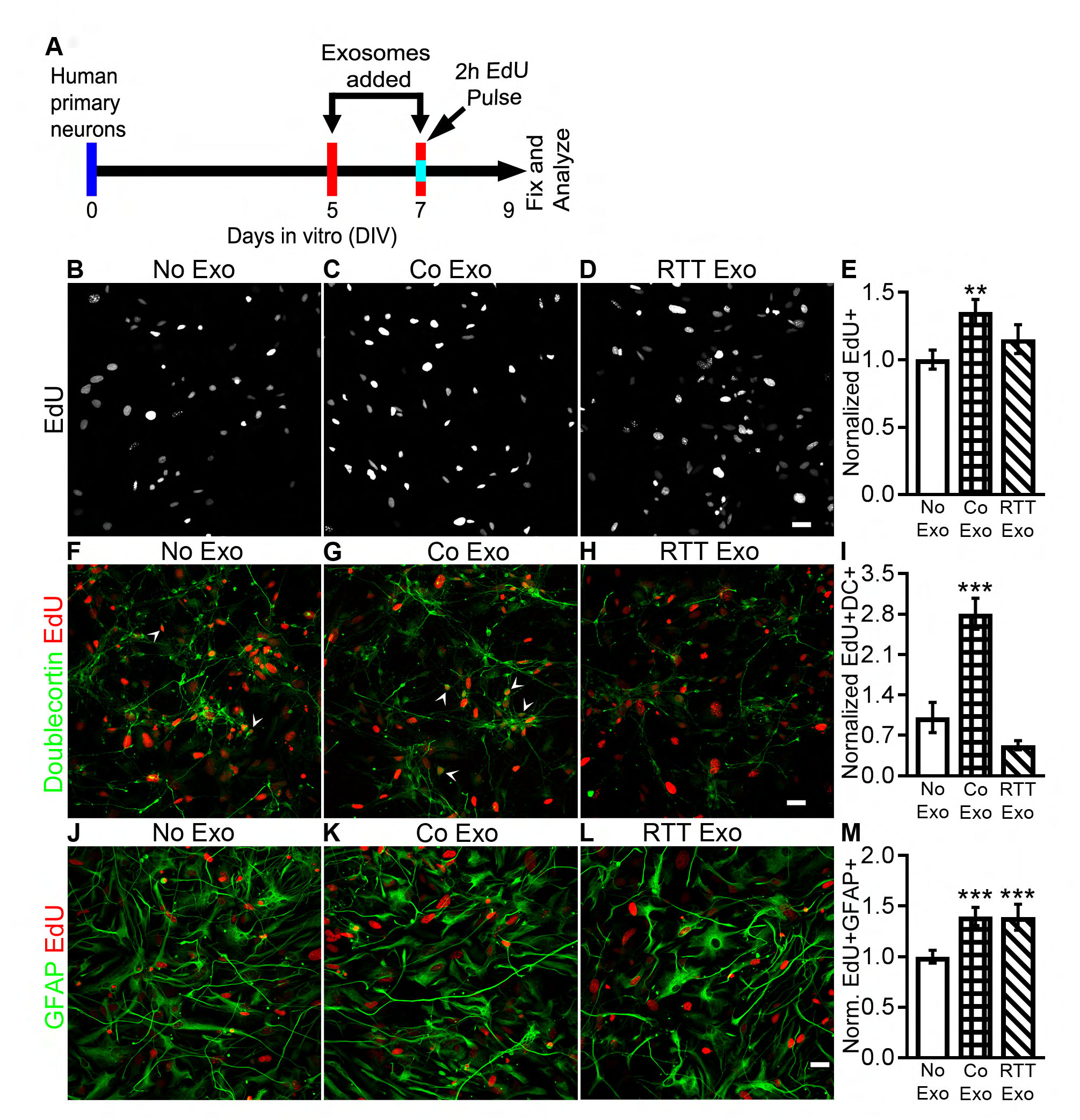
Control exosomes increase proliferation and neuronal fate in developing neural cultures while RTT exosomes have no effect. A. Protocol for treatment of human primary neural cultures with exosomes to assay proliferation, survival and cell fate. Human primary neural cultures were treated with media (No exosomes), control exosomes or RTT exosomes on DIV 5 and 7. On DIV 7, cultures were exposed to 10 μM EdU for 2 hours, then treated with media (No Exo), control exosomes or RTT exosomes. Cultures were fixed and immunolabeled for analysis on DIV 9. B-E. Confocal images show EdU labeled human neural cultures treated with media (B), control exosomes (C) and RTT exosomes (D). Treatment with control exosomes (bars with grid pattern) increased EdU labeled progeny by 1.35 ± 0.09 (*p*=0.006) whereas RTT exosome treatment (bars with diagonal stripes) had no effect (E). F-I. Confocal images of EdU (red) and doublecortin (green) labeled cultures treated with media (F), control Exo (G) and RTT Exo (H). Control exosome treatment (bars with grid pattern) increased neuronal progeny 2.8 ± 0.27 fold (p=0.005), whereas RTT exosome treatment (bars with diagonal stripes) had no effect (I). J-M. Confocal images of EdU (red) and GFAP (green) labeled cultures treated with media (J), control Exo (K) and RTT Exo (L). control as well as RTT exosome treatment (bars with diagonal stripes) increase in GFAP labeled astroglial progeny by 1.4 ± 0.09 fold (p=0.006) and 1.4 ± 0.13 fold (p=0.009) respectively (M). N= 4 wells per group. Statistics, 2 way ANOVA with Bonferroni correction. Scale bar = 20 μm.

### Control exosomes enhance neuronal fate while RTT exosomes have no effect

To investigate whether proliferation in response to exosome treatment affects the generation of neurons, we treated human primary neural cultures with exosomes, then labeled with neuronal and non-neuronal cell markers. We treated human primary neural cultures with media alone, control exosomes, or RTT exosomes on DIV 5 and 7 and provided a 2-hour pulse of 10 μM EdU prior to addition of exosomes on Day 7 (Figure 3A). On DIV 9, the cultures were fixed and immunolabeled with EdU and doublecortin (DC, an immature neuron marker; Figure 3 F-H) to quantify the generation of neurons in EdU labeled progeny. For comparison, we also immunolabeled with EdU and GFAP (an astroglial marker; Figure 3 I-K), and quantified the generation of GFAP-expressing cells with astroglial morphology in EdU labeled progeny. Control exosome treatment increased EdU^+^DC^+^ double-labeled neurons 2.8 ± 0.27 fold (*p*=0.005) fold compared to control conditions (addition of media alone, no exosomes), whereas RTT exosome treatment did not affect the numbers of EdU^+^DC^+^ double-labeled neurons (Figure 3 F-H, L). Furthermore, control exosome treatment increased the proportion of EdU^+^DC^+^ neurons in EdU labeled compared to control conditions or treatment with RTT exosomes (Figure S3C). Control exosome treatment also increased EdU^+^GFAP^+^ double-labeled astroglia 1.4 ± 0.09 fold (p=0.006) fold compared to control, addition of media alone, conditions. In this case, the RTT exosome treatment also increased EdU^+^GFAP^+^ double-labeled astroglia by 1.4 ± 0.13 fold (p=0.009) (Figure 3 I-K, M) and the proportion of EdU^+^GFAP^+^ cells in the total population of EdU^+^ cells was not significantly different between control and RTT exosome treatments (Figure S3D), indicating that increased astroglia generation was proportional to the increase in progeny. These results show that control exosomes enhance neuronal fate and promote expansion of the neuronal pool. While RTT and control exosomes had an equivalent capacity to enhance astroglial formation, RTT exosomes lacked signals that influence neuronal fate. These results are also consistent with our proteomic data analysis predicting downstream effects on the generation of neurons (Figure 2D).

### Control exosomes rescue the reduced neuron numbers resulting from MeCP2 knockdown

Our proteomic analysis showed that control exosomes contain signaling components capable of affecting neurogenesis and circuit formation. Furthermore, control and RTT exosomes differ significantly in protein cargo that are predicted to affect proliferation, neurogenesis and synapse development and maintenance. In the experiments described above, we showed that control exosomes increase total cell number and neuronal specification in control human neural cultures, whereas treatment with RTT exosomes did not affect these features in control cultures. Mutations in the *MECP2* gene are known to produce functional deficits in some of these same phenotypes of neural circuit development (Feldman et al., 2016). We were therefore interested in testing whether control exosomes could rescue the deficits in neurogenesis and synaptic maturation and function resulting from MeCP2 loss of function.

To address this hypothesis, we knocked down MeCP2 in human neural cultures using lentivirus expressing shRNA targeted to *MECP2*, or scrambled shRNA lentivirus as a control (Marchetto et al., 2010). *MECP2* shRNA lentivirus (sh*MECP2*) was added to human primary neural cultures on DIV 3 followed by treatment with exosomes on DIV 5, 7, 9 and 11 (Figure 4 A). Our proteomic analysis of exosomes indicated that protein cargo include multiple components of diverse signaling pathways which might produce different outcomes vis a vis neural development phenotypes in recipient neurons depending on the amount, or dose, of cargo delivered by exosomes. Furthermore, it is thus far not possible to approximate the amount of exosomes or their signaling capacity in intact tissue. We therefore tested the effect of 3 doses of control exosomes on MeCP2 knockdown cultures by serially diluting the purified exosomes (1X dose) to 0.5X and 0.25X relative concentrations to investigate the relationship of dose with function. The cultures were fixed on DIV 13 and immunolabeled with Synapsin and DAPI (Figure 4B-E). Synapsin is widely expressed in neurons and was used to assay neuron number and total cell counts were determined using DAPI. We imaged using wide field fluorescence microscopy to enhance detection of Synapsin immunolabeling in cell bodies. We used insulin like growth factor-1 (IGF-1) as a positive control (Figure 4E), as it has been shown to partially restore some of the MeCP2 loss of function phenotypes by increasing expression of synaptic proteins, enhancing excitatory synaptic transmission, and restoring dendritic spine densities (Marchetto et al., 2010).

**Figure 4.**
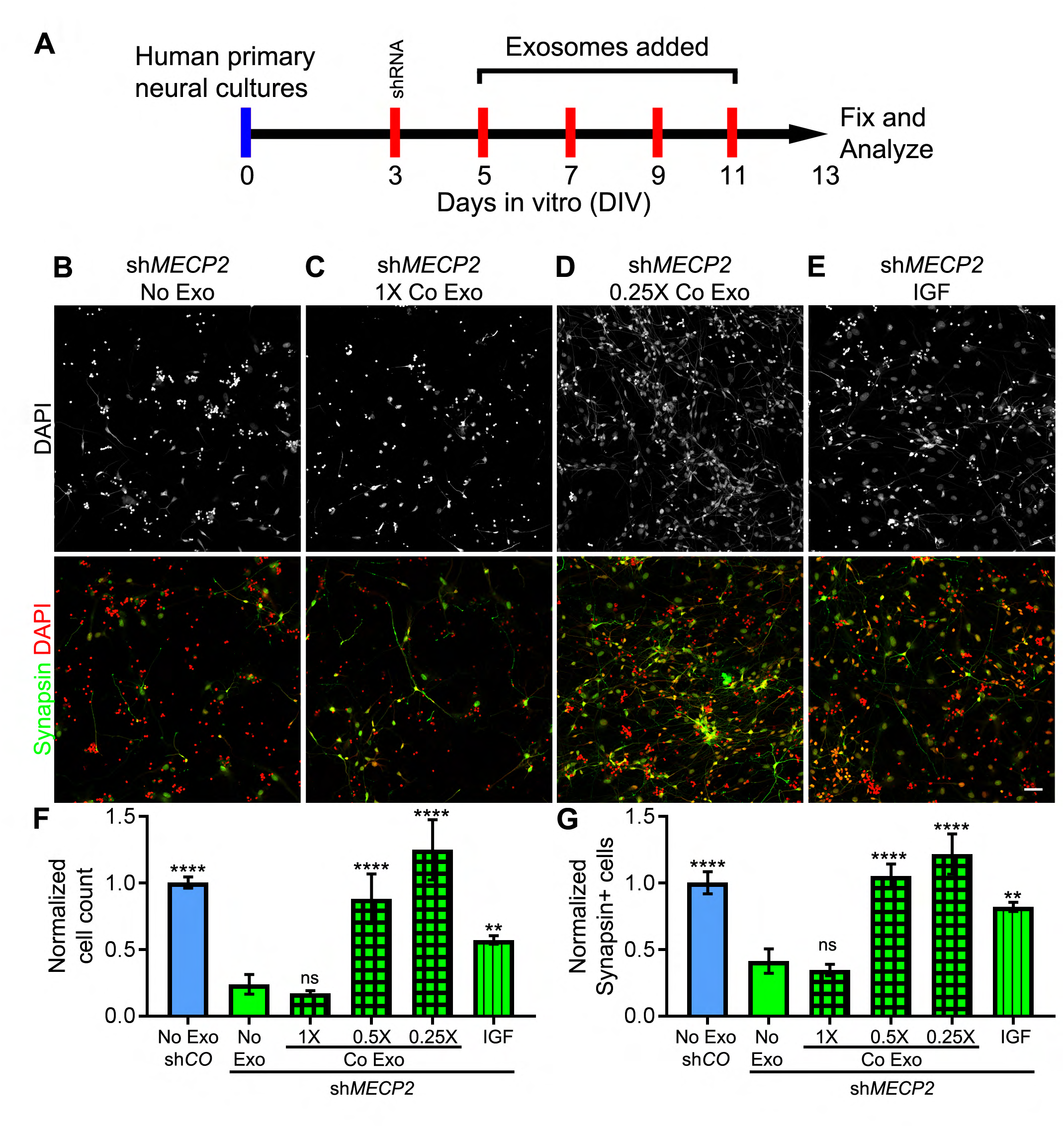
Control exosomes rescue the reduced number of neurons resulting from MeCP2 knockdown. A. Protocol for treatment of human primary neural cultures with exosomes to assay changes in number of neurons and total cells. Human primary neural cultures were infected with lentivirus expressing shRNA to knockdown MeCP2 and treated with media alone (No exosomes, open bars in F, G), 1X, 0.5X or 0.25X dose of control exosomes (bars with grid in F, G) or 10ng/ml IGF (bars with vertical stripes in F, G) on DIV 5,7,9 and 11. Lentivirus containing scrambled shRNA was used as knockdown control (shCO-No Exo). On DIV 13, cultures were fixed and immunolabeled with DAPI and the neuron marker, synapsin, to quantify the number of total cells and number of neurons, respectively. B-E. Top panels show DAPI labeling (greyscale) and bottom panels show results from an independent experiment with separate DAPI (red) and synapsin (green) double-labeled wide field fluorescent images from cultures with MeCP2 knockdown treated with media (No Exo) (B), 1X control exosomes (C), 0.25X control exosomes (D) and 10 ng/ml IGF, as a positive control. F. MeCP2 knockdown (No exosomes) reduces the number of cells to 0.24X ± 0.07 (*p*=0.0001) relative to control conditions (scrambled shRNA control: CO-No exosome). Treatment with 0.5X control exosomes and 0.25X control exosomes rescued the decreased cell number seen with MeCP2 knockdown without exosomes to 0.88 ± 0.19 (*p*=0.0001 compared to MeCP2 knockdown, no exosomes) and 1.25 ± 0.22 (*p*=0.0001 compared to MeCP2 knockdown, no exosomes) of control, respectively. IGF treatment showed partial rescue to 0.57 ± 0.03 (*p*=0.007 compared to MeCP2 knockdown, no exosomes) of control. G. MeCP2 knockdown (No exosome) decreases the number of neurons to 0.42X ± 0.09 control (CO-No exosome) (p=0.0001). Treatment with 0.5X control exosomes and 0.25X control exosomes rescued the number of neurons to 1.05 ± 0.09 and 1.22 ± 0.15 of control, respectively (p=0.0001 each compared to MeCP2 knockdown, no exosomes). IGF treatment showed partial rescue to 0.8X ± 0.03 of control (p=0.003 compared to MECP knockdown, no exosomes). N = 3 wells per group. Statistics, 2 way ANOVA with Bonferroni correction. Scale bar = 20 μm. N.S.: Not significant.

Lentiviral MeCP2 knockdown decreased total cell number and neurons compared to control lentivirus (CO-No Exo vs sh*MECP2*, No Exo; Figure 4 F, G). Treatment with control exosomes from hiPSC-derived neural cultures rescued the loss of total cell numbers (Figure 4 B-E, F) and neurons (Figure 4 B-E, G). While the 1X dose of control exosomes was ineffective, the 0.5X and 0.25X doses produced complete rescue, in contrast with partial rescue with IGF treatment. These results, together with our proteomic analysis, suggest that exosomes contain multiple signaling codes that are likely to have different outcomes depending on dose and the state of recipient cells.

### RTT exosomes have no adverse effect on total cell number or neuron numbers

Since RTT exosomes are secreted by cells that lack MeCP2 protein, they could contain negative signals that directly contribute to MeCP2 knockdown phenotypes. In this context, we were interested in testing whether RTT exosomes convey negative signals that could contribute to defects in neurogenesis and synapse development and function seen with *MECP2* mutations. Our previous results showed that two applications of RTT exosomes over 4 days did not have any negative effects on total cell number or neuronal cell fate (Figure 3 B, L), and had a positive effect on GFAP^+^ glial cell fate (Figure 3M). Here, we used a treatment protocol similar to that used in Figure 4 to test whether a10-day treatment regime with 4 applications of exosomes on DIV 5, 7, 9 and 11 has negative effects on total cell and neuronal number (Figure 5 A). Furthermore, we tested 3 doses of RTT exosomes by serially diluting the 1X stock to 0.5X and 0.25X. All cultures were infected with control lentivirus expressing scrambled shRNA to match control conditions described above and in Figure 4. The cultures were fixed on DIV 13 and the numbers of neurons and total cells were determined. Treatment with control exosomes increased total cell number (Figure 5 B-C, F) as well as the number of neurons compared to addition of media alone (Figure 5 B-C, G). Treatment with RTT exosomes did not show any adverse effect on total cell number (Figure 5 D-E, F) or numbers of neurons (Figure 5 D-E, G). On the contrary, the 0.5X dose of RTT exosomes increased the total cell number compared to control cultures without exosome addition (Figure 5 F). These results suggest that RTT exosomes do not have a dominant acute deleterious effect on cell proliferation or neuronal differentiation.

**Figure 5.**
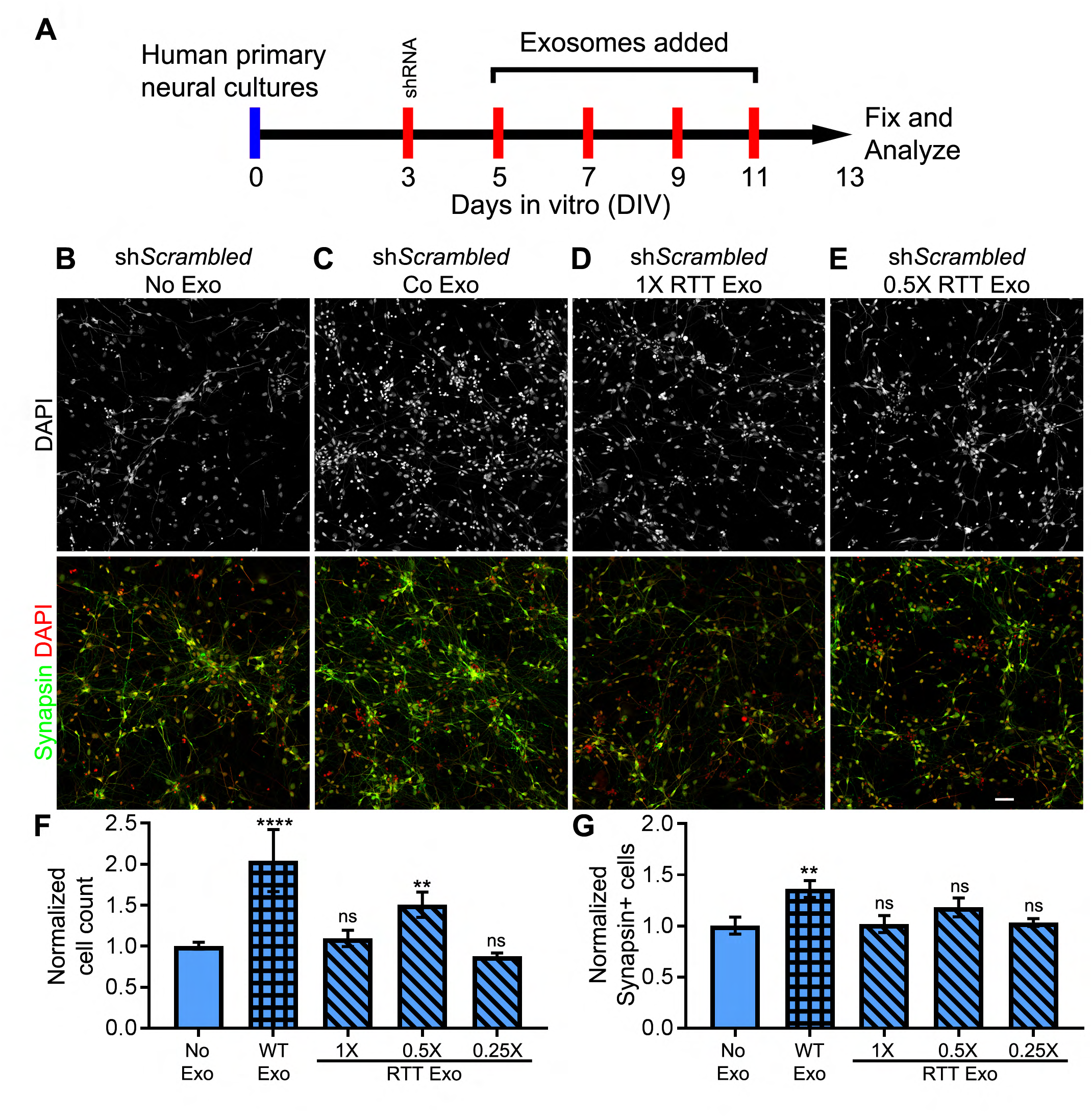
RTT exosomes have no adverse impact on total cell number or neuronal number. A. Protocol for treatment of human primary neuron cultures with exosomes to assay change in number of neurons and total cells. Human primary neurons were infected with lentivirus expressing scrambled shRNA and treated with media (No exosomes, open bars in F, G), control exosomes (bars with grid pattern), 1X, 0.5X or 0.25X doses of RTT exosomes (bars with diagonal stripes) on DIV 5,7,9 and 11. On DIV 13, cultures were fixed and immunolabeled with DAPI and differentiated neuron marker synapsin to quantify number of total cells and number of neurons. B-E. Wide field fluorescence images show DAPI (greyscale, red) and synapsin (green) labeled cultures treated with media (B), control exosomes (C), 1X RTT exosomes (D) and 0.25X RTT exosomes (E). Data shown in top and bottom panels are examples from separate experiments. F. Control exosome treatment lead to 2.05X ± 0.69 (p=4.4×10^−4^) increase in number of cells compared to control (No exosome). The treatment with 0.5X RTT exosomes increased the number of cells by 1.48X ± 0.38 (p=0.0001). 1X RTT and 0.25X exosome treatment were not statistically different from control. G. Control exosome treatment increases the number of neurons 1.36X ± 0.08 (*p*=0.004) compared to control (No exosomes). None of the RTT exosomes treatments were statistically different from control. N=3 wells for each group. Statistics, 2 way ANOVA with Bonferroni correction. Scale bar = 20 μm. N.S.: Not significant.

### Control exosome treatment rescues increase in cell death upon MeCP2 knockdown

The results presented above (Figures 4, 5) indicate that treatment of MeCP2 knockdown cultures with control exosomes and treatment of control knockdown cultures with RTT exosomes affect total cell number and neuron numbers in a dose dependent manner. Here, we tested whether exosome treatments affect cell survival. To this end, we compared the effects of control and RTT exosome treatments on human neural cultures with MeCP2 knockdown or control cultures, infected with scrambled shRNA lentivirus, following the treatment protocol as previously shown (Figures 4 and 5). We assayed apoptotic cells, quantified using DAPI staining of nuclear morphology. Treating control cultures with RTT exosomes did not increase the proportion of apoptotic cells (Figure 6A-C). In addition, in control cultures, treatment with control exosomes or 3 doses of RTT exosomes, did not affect the number of apoptotic cells across treatment conditions, compared to control cultures without exosomes (Figure 6SA). These results suggest that RTT exosomes do not confer toxicity to control recipient neural cultures.

**Figure 6.**
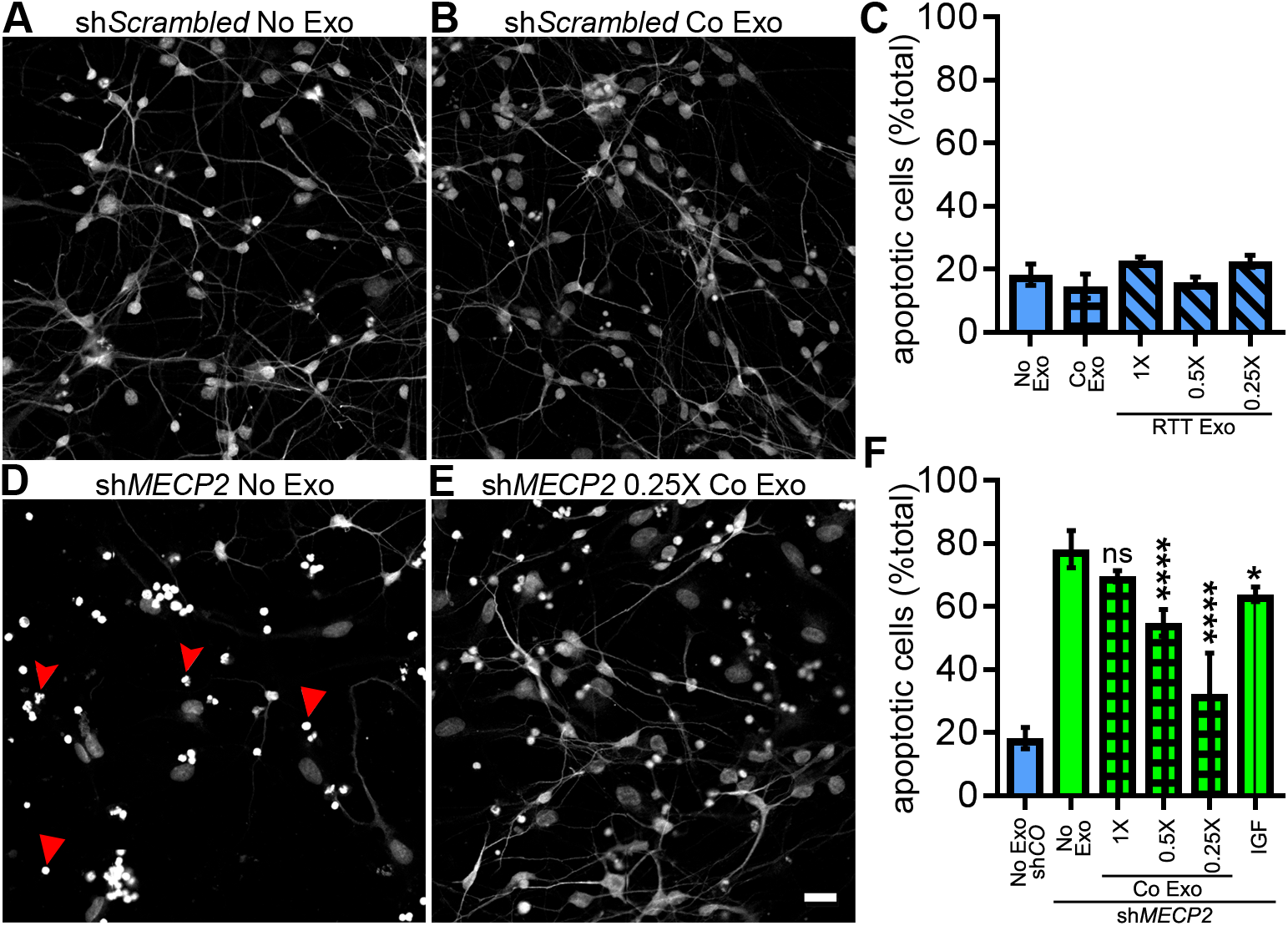
Control exosomes rescue increased cell death resulting from MeCP2 knockdown. A-C. RTT exosomes do not affect apoptosis. Cultures were infected with lentivirus expressing scrambled control shRNA (shScrambled) and treated with media without exosomes, control exosomes or with RTT exosomes (1X, 0.5X or 0.25X doses), as described in Figure 5A. A, B. Wide field fluorescence images of cultures treated with media (A, no Exo) and with control exosomes (B). C. Quantitative analysis of the fraction of cells that were apoptotic in cultures treated with control shSrambled lentivirus. Treatments with control exosomes (bars with grids) or RTT exosomes (bars with diagonal stripes) does not increase the proportion of apoptotic cells compared to media control. D-F. Control exosomes rescue the increased apoptosis seen with MeCP2 knockdown. Cultures were infected with lentivirus expressing shRNA for *MECP2* (sh*MECP2*) and treated with media without exosomes, or control exosomes (1X, 0.5X or 0.25X doses) or IGF as described in Figure 4. D, E. Wide field fluorescence images of cultures treated with media (D) and 0.25X control exosomes (E). Arrowheads show apoptotic cells (D). F. Quantitative analysis of the fraction of cells that were apoptotic in MeCP2 knockdown cultures compared to cultures treated with control shSrambled lentivirus (blue bar). MeCP2 knockdown increased the fraction of apoptotic cells 4.2 fold, compared to control, from 18.2 ± 3.4% to 78.3 ± 5.8% of total cells (*p*=0.0001) (sh*MECP2*, no exosome, solid green). Treatments with control exosomes (0.5X and 0.25X doses) significantly decreased reduced the fraction of total cells that were apoptotic, from 78.3 ± 5.8% to 54.9. ± 4.3% (p=0.0001) and 32.3 ± 12.8% (*p*=0.0001) respectively. IGF treatment (10ng/ml) decreased apoptotic cells to 64.0 ± 2.3% of the total cell numbers (*p*=0.01). N=3 wells for each group. Statistics, 2 way ANOVA with Bonferroni correction. Scale bar = 20 μm.

To test whether control exosome treatments affect apoptosis in MeCP2 loss of function conditions, cultures were infected with *MECP2* shRNA lentivirus on DIV 3 followed by treatment with control exosomes on DIV 5, 7, 9 and 11. MeCP2 knockdown without added exosomes (No exo) significantly increased the fraction of apoptotic cells by 4 fold (Figure 6 D-F) and increased the relative number of apoptotic cells by about 3.5 fold (Figure 6SB) compared to control conditions (shCO-No exo). Comparing the fraction of apoptotic cells within each treatment condition indicates that the fraction of apoptotic cells seen with MeCP2 knockdown was significantly reduced by treatment with 0.5X and 0.25X control exosomes (Figure 6 D-F). While IGF treatment significantly reduced the fraction of apoptotic cells (Figure 6F), it did not change the number of apoptotic cells (Figure 6SB). While treatment with the 1X dose of control exosomes significantly decreased apoptosis in MeCP2 knockdown cultures compared to cultures without exosome treatment, the 0.5X and 0.25X doses did not rescue the increased apoptosis in MeCP2 knockdown cultures (Figure 6SB). The effect of different control exosome doses on the fraction of apoptotic cells (Figure 6 F) reflects a non-linear relationship compared to the effect of different control exosome dose treatments on the increase in number of neurons and total cells, as shown in Figure 4F, G. It is interesting to note that while treatment with the 1X control exosome dose rescued the number of apoptotic cells in MeCP2 knockdown cultures, it failed to rescue the decrease in number of neurons and total cell phenotype of MeCP2 knockdown, also seen in Figure 4. The observation that control exosome dose responses are different for mediating rescue of different phenotypes, specifically the total cell and neuron numbers compared to apoptotic cells, supports our prediction from the proteomic data that exosomes contain complex signaling information affecting different biological outcomes.

### Control exosome treatment increases synaptogenesis and synchronized firing in RTT iPSC-derived neurons

RTT hiPSC-derived neuronal cultures have fewer synapses and less activity-driven calcium transients than control neurons (Marchetto et al., 2010). We tested whether control exosomes have the capacity to rescue these synaptic and circuit deficits in RTT cultures. To assay the effect of exosome treatment on synaptogenesis, we differentiated RTT hiPSC-derived NPCs into mature 6 week-old neural cultures and the cultures were treated 4 times with 1X or 0.25X doses of control exosomes over 8 days, as described in Figure 7A. After exosome treatments, the cultures were fixed and labeled with antibodies against MAP2 to label neurons and synapsin I to label presynaptic puncta (Figure 7B,C). Images were analyzed for quantitative changes in synaptic puncta number and intensity. Neural cultures treated with the 0.25X dose of control exosomes had significantly increased puncta density compared to cultures without exosome treatment, whereas treatment with the 1X dose of exosomes resulted in an intermediate puncta density (Figure 7 B-D). Furthermore, treatments with control exosomes shifts the distribution of puncta intensities toward lower intensities (Figure 7E), suggesting that the increase in synaptic density seen in Figure 7D (change to F with update in panel labeling) is due to an increase in lower intensity puncta. Together, these data suggest that control exosomes increase synaptogenesis and circuit development in RTT neural cultures.

**Figure 7.**
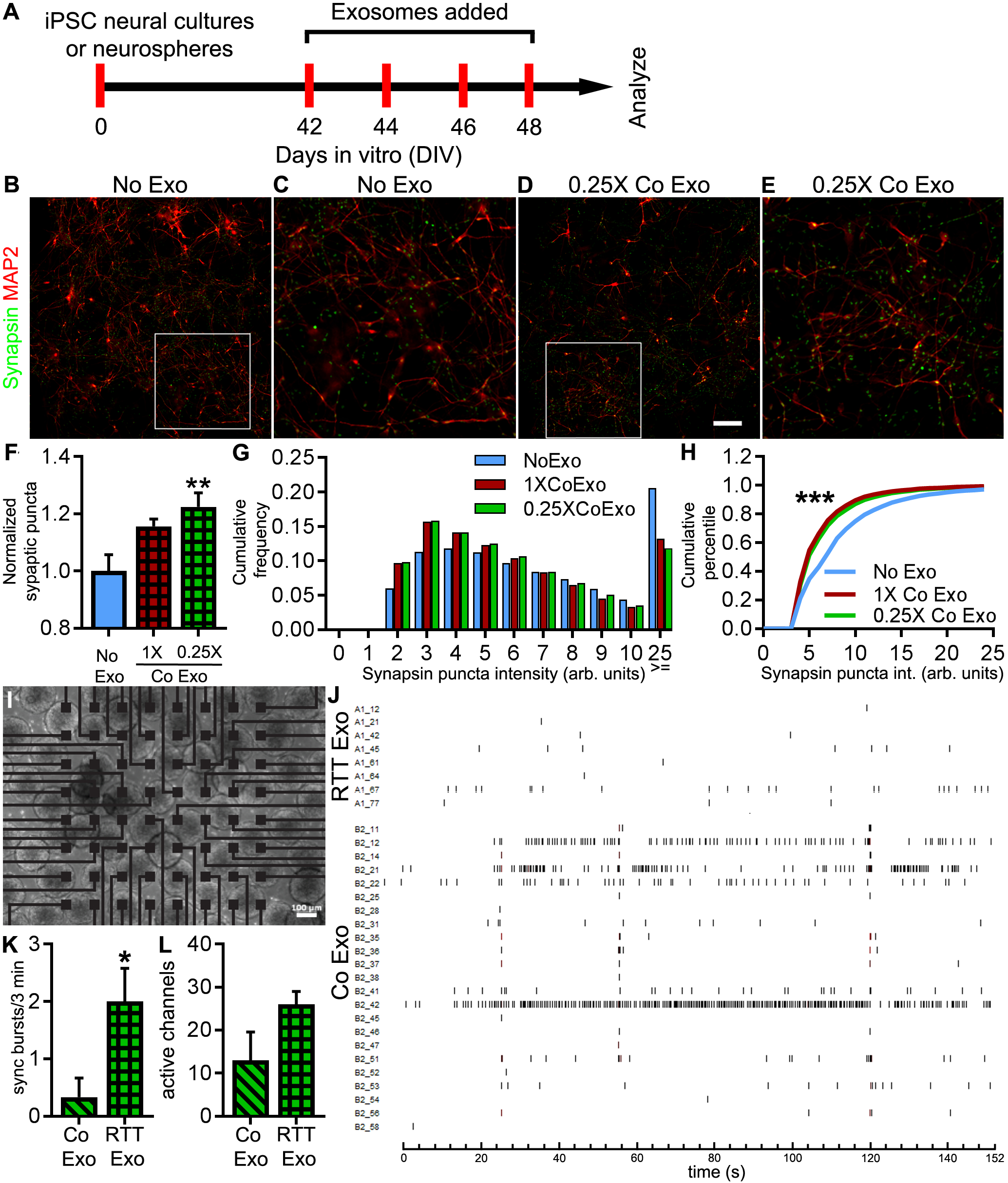
Control exosome treatment increases synaptogenesis and synchronized firing in RTT iPSC-derived neurons. A. Protocol for treatment of RTT iPSC-derived neural cultures or neurospheres with control exosomes to assay synaptic puncta or synchronized firing. B-H. Control exosomes increase synaptogenesis in RTT iPSC-derived neural cultures. Cultures were fixed on DIV 50 and synaptic puncta density and intensity were quantified. Images of MAP2 (red) and synapsin (green,) labeling in RTT neural cultures without exosome treatment (B, C) or with control exosome treatment (D, E). C, E. Enlarged regions marked by inset boxes in B and D, respectively. F. Exosome treatments increased synapse density. Treatment with control exosomes (0.25X dose)
increased presynaptic puncta density to 122X ± .05 of no exosome treatment control values (*p*=0.006; n=4 wells each, 2 way ANOVA with Bonferroni correction). G. The frequency distribution of synaptic puncta intensity shows an increase in low intensity puncta with control exosome treatments (blue bar, no exosomes; red bar, 1X control exosomes; green bar, 0.25X exosomes). H. Kolmogorov-Smirnoff plot of cumulative frequency of synaptic puncta intensity shows that control exosome treatments (1X control (red line, D-stat=0.24, p value <0.0001) and 0.25X control (green line, D-stat=0.21, p value <0.0001)) increase the fraction of low intensity puncta compared to no exosome treatment (blue line). I-L. Treatment of RTT neurospheres with control exosomes increases neural circuit activity. RTT hiPSC-derived neurospheres were plated on 64-channel multi-electrode array (MEA; I) and treated with RTT exosomes or control exosomes. J. Aligned raster plots of spiking activity over 3 minutes of recordings from independent active channels show less overall activity and less synchronized activity in neurospheres treated with RTT exosomes compared to those treated with control exosomes. K. Graph showing synchronized bursts of activity occur with a greater frequency in neurospheres treated with control exosomes compared to neurospheres treated with RTT exosomes (RTT exo: 0.33 ± 0.33 bursts/3 min, control exo: 2.0 ± 0.68 bursts/3min (*p*=0.03, n= 3 arrays each, two tailed t test). L.Graph showing the number of active channels is not significantly different in neurospheres treated with control exosomes compared to RTT exosomes (RTT: 13 ± 6.56, control: 26 ± 3.0). Scale bar = 50 μm (B, D), 100 μm (F).

Neurospheres generated from hiPSC-derived neural cells are a convenient 3dimensional experimental system for measuring neuronal activity and connectivity in a network using Multi Electrode Arrays (MEAs). Neurospheres mature into a network that displays synchronized neuronal firing (Nageshappa et al., 2016). To analyze whether control exosome treatments affect circuit development, we assayed synchronized neuronal firing in neurospheres. We generated neurospheres from hiPSC RTT-derived NPCs and plated them onto MEAs (Figure 7H). Neurospheres were treated with exosomes 4 times over 8 days, as described (Figure 7A) and activity across the MEA was evaluated. Raster plots of spike recordings from individual electrodes shows that RTT neurospheres treated with RTT exosomes were less active than neurospheres treated with control exosomes. In particular, neurospheres treated with control exosomes had more activity per channel and more active channels than neurospheres treated with RTT exosomes (Figure 7 G, H). The aligned raster plots of spiking activity from independent active channels show that neurospheres treated with control exosomes had significantly more synchronized activity, measured by synchronous bursts of spikes across multiple channels (Figure 7 G, I). The increase in synchronous activity extending across multiple electrodes with control exosome treatment indicates that exosome treatment increases network connectivity. Together, these results show that exosome treatment is not only capable increasing synaptic connections, but exosomes also enhance coordinated activity in neural circuits.

## Discussion

The potential function of exosomes in neural circuit development is unclear. Here we show that exosomes play a profound role in neural circuit development leading to enhanced proliferation, neuronal specification and circuit connectivity. Furthermore, we show that exosomes from RTT patient hiPSC-derived neural cultures, a model of RTT, fail to promote neural circuit development, while exosomes from isogenic control hiPSC-derived neural cultures rescue developmental deficits, including neurogenesis, synaptogenesis and circuit connectivity, in human neural cultures or neurospheres with MeCP2 knockdown. Quantitative proteomic analysis showed that exosomes from human neural cultures are enriched in proteins known to play diverse roles in neuronal circuit development. Exosomes from RTT and control neural cultures differed significantly in signaling proteins involved in neuronal circuit development. Together our data demonstrate that exosomes facilitate the development of human neural circuits and that treatments with control exosomes rescue neurodevelopmental defects in a model of RTT.

### Proteomic analysis of exosomes

Analyses of the proteomic content of exosomes from glioblastoma cells and neurological diseases of the aging brain, such as Parkinson’s Disease, have led to several proposed functions of exosomes, such as delivery of signaling components that might modify the local environment and facilitate tumor growth or trans-cellular transport of toxic proteins, in the case of proteinopathies (Basso and Bonetto, 2016; Chiasserini et al., 2014; Jayaram et al., 2014; Tomlinson et al., 2015). Recent work suggests that the EphB/ ephrinB axon guidance molecules are present in exosomes and can regulate growth cone collapse in vitro (Gong et al., 2016), and that intercellular signaling by exosomes may play a role in synapse elimination (Bahrini et al., 2015). However, no study to our knowledge has shown any analysis of exosomes in the context of normal neural development or neurodevelopmental disorders. We now provide quantitative analysis of differences in proteomic content of exosomes from RTT syndrome patient iPSC-derived neural cultures with MeCP2 loss of function compared to CRISPR/Cas9-corrected isogenic control exosomes. Neural cultures from patient-derived iPSCs offer several advantages to model aspects of neurodevelopmental disorders, including RTT syndrome and other autism spectrum disorders (Beltrao-Braga and Muotri, 2017; Freitas et al., 2014; Sztainberg and Zoghbi, 2016). Here, hiPSC-derived neural cultures were particularly valuable for the generation of large volumes of conditioned media necessary for exosome purification. The use of isogenic control neural cultures generated by correcting the *MECP2* mutation minimizes effects due to differences in genetic background to parse out differences specifically resulting from *MECP2* mutation. Our proteomic analysis identified ∼240 differentially-expressed neural proteins between RTT and control exosomes, which are predicted to affect proliferation, neuronal development and synaptic maturation of neurons, leading to abnormal connectivity of neural circuits. The proteins in our differentially expressed dataset predicted to affect proliferation include Plexin B1/B2 (Daviaud et al., 2016; Saha et al., 2012), Ephrin B3 (Ricard et al., 2006), delta/notch like EGF repeat containing (DNER) (Hsieh et al., 2013), and neurotrophic receptor tyrosine kinase 2 (NTRK2) or Trk-B (Bartkowska et al., 2007). The proteins affecting neurogenesis or neuronal development include cell division cycle 42 (CDC42) (Cappello et al., 2006), Cadherin 2 (CDH2) or N-Cadherin (Yagita et al., 2009), Ephrin B1 (EFNB1) (Klein and Kania, 2014; Qiu et al., 2008), GDNF family receptor alpha 1 (GFRA1) (Marks et al., 2012) and Trk-B (Li et al., 2008). The proteins affecting synaptic maturation include Solute carrier family 12 member 2 (SLC12A2) or NKCC1 (Pfeffer et al., 2009), Adhesion G protein-coupled receptor L3 (ADGRL3) or Latrophilin-3 (O’Sullivan et al., 2012), Fibronectin leucine rich transmembrane protein 1 (FLRT1) (O’Sullivan et al., 2012), Neuroligin-2 (NLGN2) (Chih et al., 2005; Scheiffele et al., 2000), and Contactin 1 (CNTN1) (Puzzo et al., 2013). It is important to note that for a given function, both positive as well as negative regulators are present in exosomes. In addition, many class of signaling molecules like Ephrins and Trk receptors, are widely implicated in neuronal development and function. Taken together, these data suggests that the effect of exosome signaling is likely to be a combinatorial sum of overlapping and sometimes opposing signaling messages. This is consistent with our observation that different doses of exosomes generate different responses in proliferation compared to apoptosis.

### Exosome function in neuronal circuit development

Our data indicate that exosomes play multiple roles in neuronal circuit development. Given the spatial and temporal complexity of circuit development, we used an experimental design, in which we examined the effects of exosome treatments at different developmental time points of circuit development. This allowed us to demonstrate effects of exosomes on cell proliferation, assayed at relatively early time points, neuronal differentiation and apoptosis, assayed at intermediate time points, and synaptogenesis and circuit connectivity, assayed at later time points. These diverse outcomes of exosome treatments are consistent with proteomic cargo identified in control exosomes.

### Control exosomes rescue neurodevelopmental deficits of MeCP2 knockdown

Our data indicate that treatment with exosomes released from control neural cultures rescue several phenotypes of deficient neural circuit development seen with MeCP2 knockdown. For instance, control exosomes rescue deficits in neurogenesis, synaptogenesis and neural circuit connectivity in MeCP2 knockdown cultures. These data are consistent with the proteomic data indicating that exosome cargo are predicted to affect multiple processes that occur over the course of neural development. An important feature of exosomes is that they can signal over a variety of length scales and act non-cell autonomously. This may be particularly relevant to studies that have shown that restoration of MeCP2 function in glia can rescue neuronal and behavioral deficits in mouse models of RTT, consistent with long range non-cell autonomous mechanisms underlying rescue (Lioy et al., 2011). We evaluated the effects of exosomes released from mixed neural cultures, including both neurons and astrocytes, suggesting that the outcomes we identify may arise from signaling from multiple cell types.

Rett syndrome displays a complex relationship with between MeCP2 loss of function and clinical outcome (Feldman et al., 2016; Leonard et al., 2017; Zoghbi, 2016). Environmental factors and X-linked inactivation are important causes of variations in clinical outcome (Leonard et al., 2017; Vacca et al., 2016; Zoghbi, 2016). Specifically, X-linked inactivation leads to mosaicism where some fraction of cells are wildtype for MeCP2. Our results showing that exosomes from control cells rescue MeCP2 loss of function phenotypes in mutant cells offer a possible mechanistic explanation of the phenotypic variation. If a sufficient number of wild type cells are present, they could signal via exosomes to rescue the mutant cells, providing an additional factor for explanation of milder clinical outcomes in some patients.

Exosomes have been hypothesized to play a pathological role in several neurological disorders, particularly proteinopathies, by spreading pathological molecules to healthy tissue. Alternatively, exosomes have been postulated to play a protective role by dumping pathological molecules out of cells (Howitt and Hill, 2016; Quek and Hill, 2017). In the context of healthy human neural cultures, we find that exosome treatment promotes neural development. In the context of the neurodevelopmental deficits resulting from MeCP2 loss of function, our data do not support a protective or pathogenic role for exosomes. Treating human primary neurons with exosomes from RTT patient iPSC-derived neural cultures does not affect have a negative impact on proliferation or neuronal cell fate specification, nor an effect on apoptosis. These data show that RTT exosomes do not convey signals that disrupt neurogenesis or development of neuronal connectivity in control cultures, indicating that they do not play a pathogenic role in circuit development. Furthermore, RTT exosomes neither rescue nor exacerbate neuronal circuit defects resulting from MeCP2 knockdown. Our proteomic data suggest that the differences in the content of RTT and control exosomes may reflect the disruption of biological functions known to occur with MeCP2 loss of function, consistent with exosomes serving as biomarkers of the dysfunctional state of the cells with *MECP2* mutation.

In conclusion, our data indicate for the first time that hiPSC-derived neural exosomes have a functional impact in a neurodevelopmental disease model. More importantly, we show that exosomes are able to reverse some of the pathological phenotypes observed in RTT neurons, indicating that hiPSC-derived exosomes could have a wide range of new therapeutic applications in neurodevelopmental disorders in the future.

## Acknowledgments

We thank Kristin Baldwin and Ardem Patapoutian from comments on the manuscript and members of the Cline lab for helpful discussions. This work was supported by grants from the National Institutes of Health (R01MH108528, R01MH094753, R01MH109885, R01MH100175, U19MH107367), a SFARI grant #345469, and a NARSAD Independent Investigator Grant to A.R.M, an International Rett syndrome foundation (IRSF) fellowship to P.M., NIH grants (R01MH103134 and R01EY011261) and an endowment from the Hahn Family Foundation to HTC and a Fellowship from the California Institute of Regenerative Medicine (CIRM, TG2-01165) to PS, NIH grants 5R01MH067880 and 5 R01 MH100175 to JRY. The authors declare no competing interests.

## Author contributions

Conceptualization, PS and HTC; Methodology, PS, CC, PM, LS, DM; Investigation, PS, PM, CC, DM; Visualization: PS, Writing – Original Draft, PS and HTC; Writing – Review, PS, HTC, PM, CC, LS, DM, JY, AM; Funding Acquisition, PS, PM, HTC, AM, JRY.

---

**Figure 2S:**
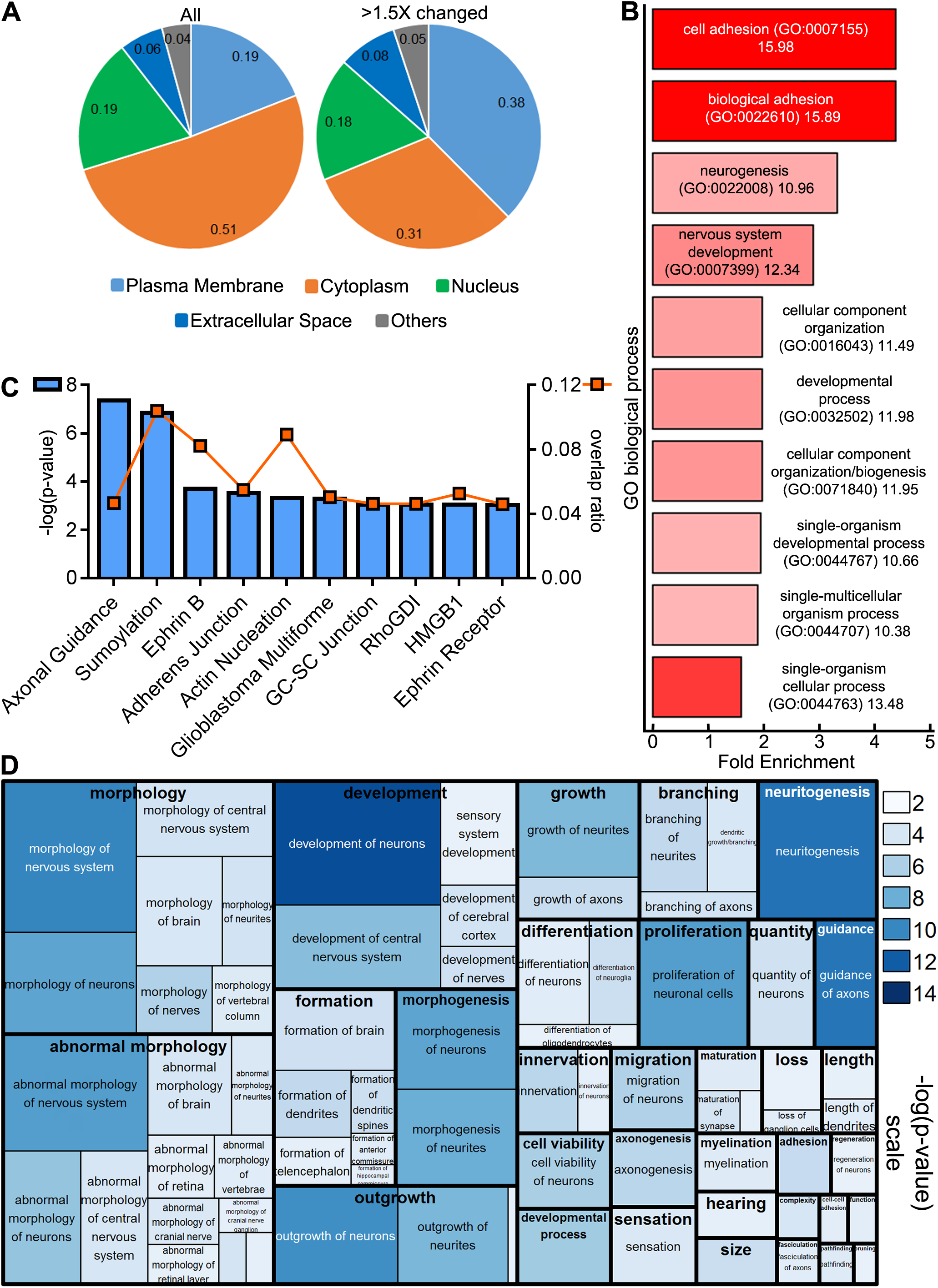
Analysis of exosome proteomics: Comparison between experimental replicates and identification of complex signaling components that are altered with MeCP2 mutation. Related to figure 2. A. Comparative analysis of the top 10 canonical pathways from Ingenuity analysis using the data from two experimental replicates, experiment 1 (blue bars, blue squares) and 2 (orange bars, orange circles). Bars, plotted on the left Y axis show significance values as -log(p-value). The line charts with symbol markers plotted on the right Y axis show overlap ratios of the number of proteins in our experimental dataset versus the total number of proteins annotated by IPA to each canonical pathway. B. Comparative Ingenuity analysis of the relative strength of the categories “downstream effects for biological functions” and “diseases in the nervous system” for the two experimental replicates, experiments 1 and 2. The significance values are plotted as -log(p-value) heatmap ranging from 0, white to 80, red, with -log(p-value) for each disease and bio-function specified in the box. C. Top 10 canonical pathways from Ingenuity analysis using the combined dataset of 2572 proteins common to both experiments. Blue bars, plotted on the left Y axis show significance values as -log(p-value) and the orange square markers, plotted on the right Y axis show the overlap ratios of the number of proteins from our dataset relative to total number of proteins annotated to each canonical pathway. D. Treemap heatmap showing Ingenuity analysis of the relative strength of downstream effects for biological functions and diseases in the nervous system using the combined dataset of 2572 proteins. The predicted biological functions are plotted hierarchically as a treemap, where the heatmap with the color look up table indicates the significance values plotted as -log(p-values). The box area represents the number of proteins from the dataset annotated for each given function ranging from 34 (morphogenesis of axons) to 306 (development of neurons).

**Figure 3S:**
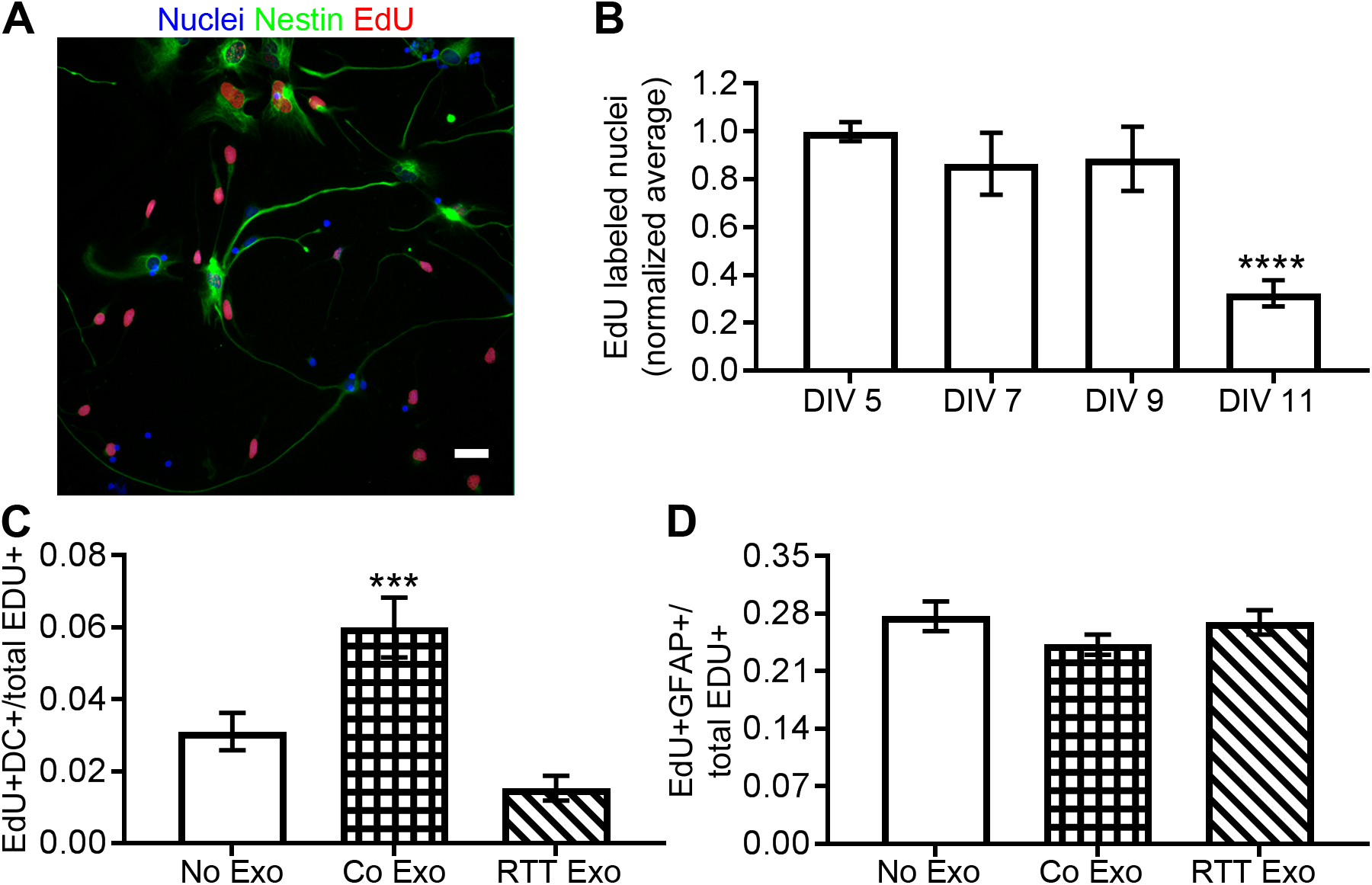
Primary human neural cultures include progenitor cells. Control exosome treatment increases neurogenesis. Related to figure 3. A. Immunolabelling of human primary neural cultures with Nestin (green) showing neural progenitor cells (NPCs) that also co-label with EdU (red). B. To test whether human primary neural cultures maintain proliferative activity over 11 DIV, cultures were treated with EdU for 4 hours on DIV 5, 7, 9 and 11 and immediately fixed. Cultures were immunolabeled for EdU and counterstained with DAPI and analyzed to determine for the number of EdU immunolabeled cells, normalized to DAPI+ nuclei. Data for each timepoint were then normalized to 5 DIV. EdU incorporation in human primary neural cultures was constant over 5-9 DIV and then dropped significantly to 32.3 ± 5.5% compared to DIV 5 (*p*=0.0001, 2 way ANOVA with Bonferroni correction) at DIV 11. N= 4 wells each. C-D. Human primary neural cultures were treated with media (No exosomes, blank bar), control exosomes (bar with grid pattern) or RTT exosomes (bars with diagonal stripes pattern) on DIV 5 and 7 as described in Figure 3A. On DIV 7, cultures were exposed to 10 m at DIV 11. N= 4 wells each. C-D. Human primary neural cultures were treated with media (No exosomes, blank bar), control exosomes (bar with grid patter for analysis of neuronal specification. Control exosome treatment doubled the fraction of DC+ neurons among newly synthesized progeny from 3.1 ± 0.5% to 6.0 ± 0.8% (p=0.006) whereas the fraction of newly generated neurons after RTT exosome treatment was not different from control cultures treated with media (C). The fraction of GFAP^+^ cells among newly synthesized progeny was unchanged with exosome treatments (D). N=4 wells per group. Scale bar = 20 μm.

**Figure 4S:**
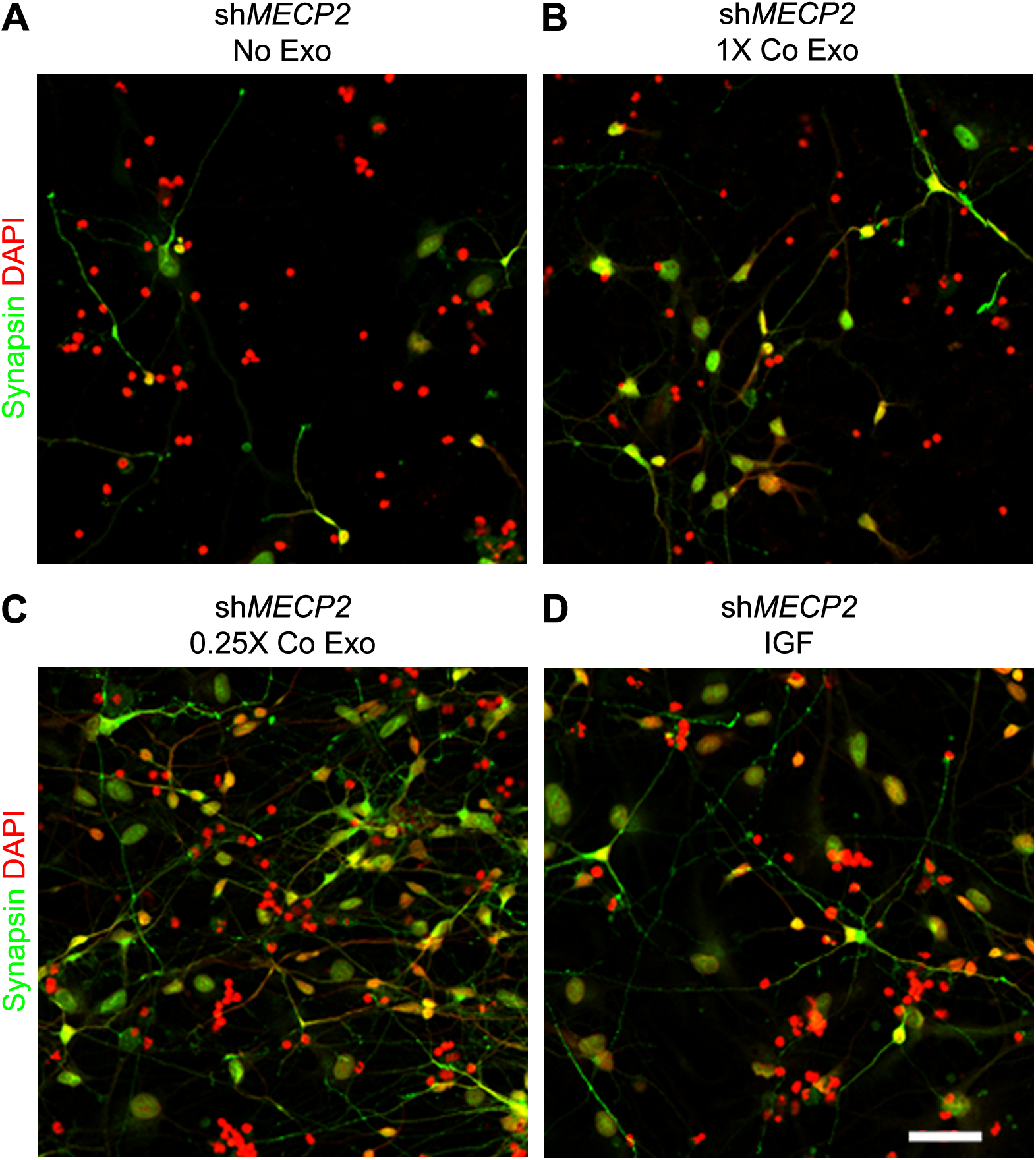
Control exosomes rescue the reduced number of neurons resulting from MeCP2 knockdown. Related to figure 4. Wide field fluorescence images of DAPI (red) and synapsin (green) labeled cultures with MeCP2 shRNA knockdown treated with media (No Exo, A), 1X control exosomes (1X Co Exo, B), 0.25X control exosomes (0.25X Co Exo, C) and 10 ng/ml IGF (D) as described in Figure 4A. The images show 3.5X zoom of images in Figure 4B-E bottom panel. N=10 fields for each group. Scale bar = 20 μm.

**Figure 5S:**
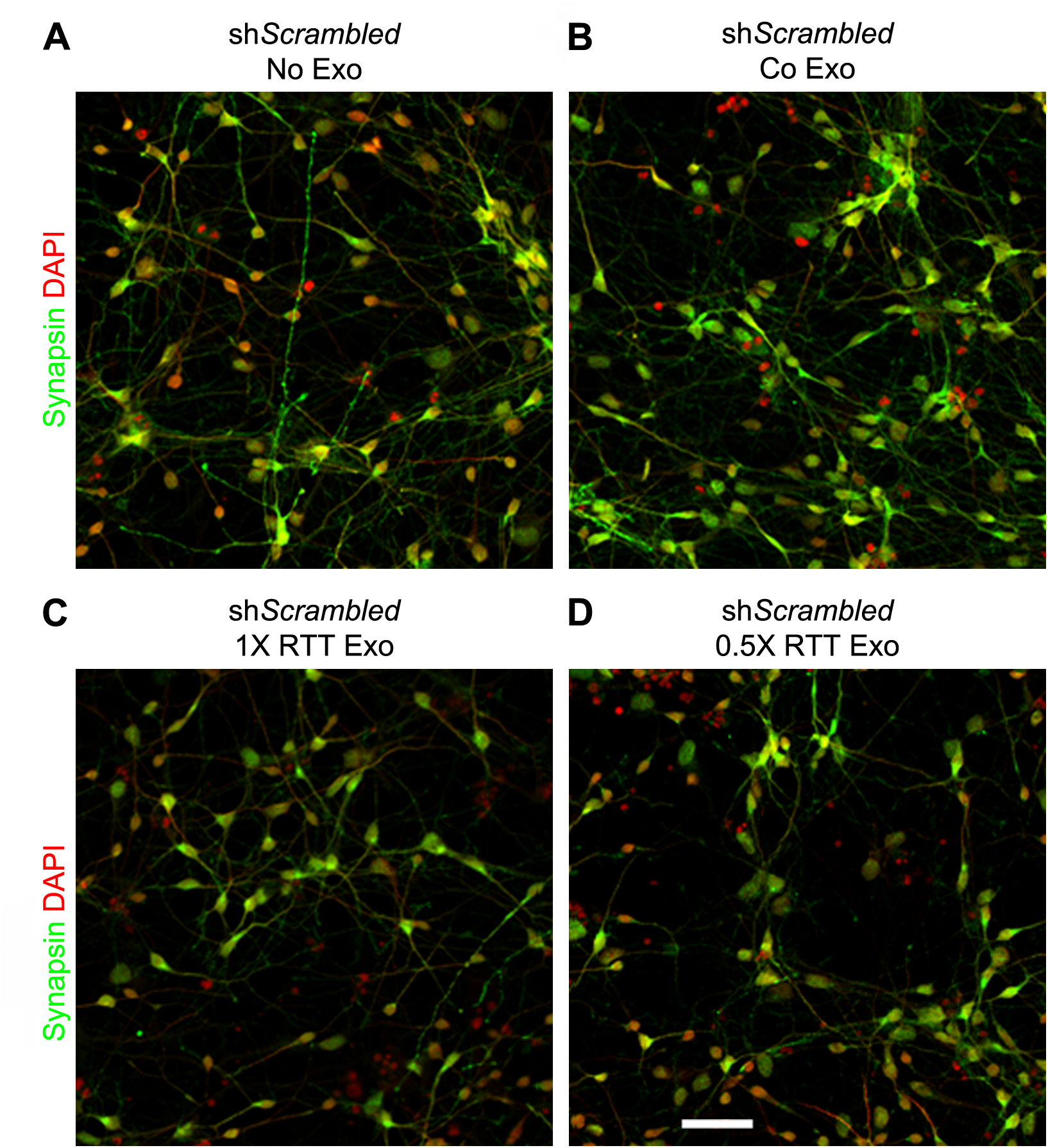
RTT exosomes had no adverse impact on total cell number or neuronal number. Related to figure 5. Wide field fluorescence images show DAPI (red) and synapsin (green) labeled cultures with scrambled shRNA control treated with media (No Exo, A), control exosomes (Co Exo, B), 1X RTT exosomes (C) and 0.5X RTT exosomes (D) as described in Figure 5A. The images represent 3.5X zoom of images in Figure 5B-E bottom panel. N=10 fields analyzed for each group. Scale bar = 20 μm.

**Figure 6S.**
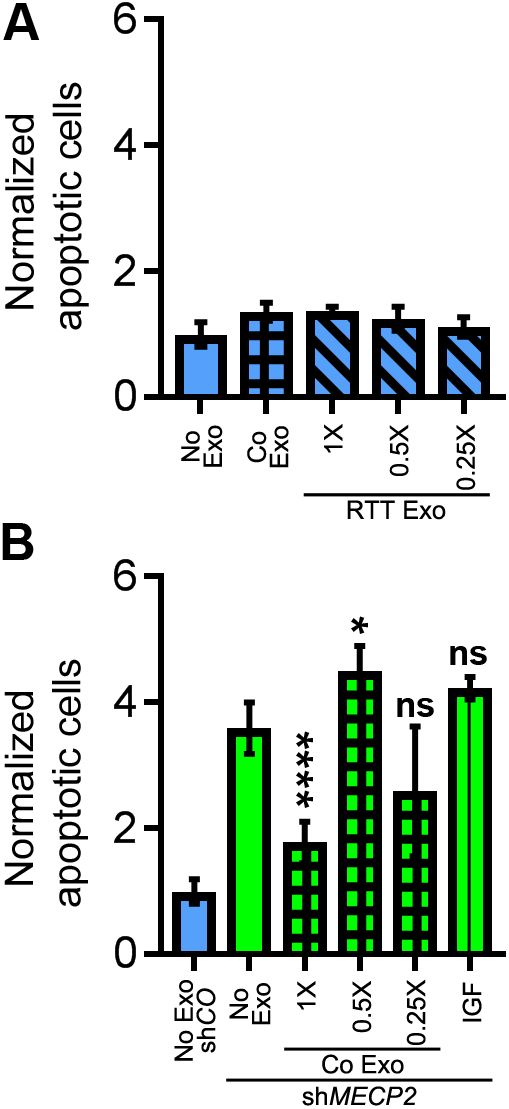
Control exosome treatment rescues increase in cell death upon MeCP2 knockdown. Related to figure 6. A. RTT exosomes do not affect apoptosis. Cultures were infected with lentivirus expressing scrambled control shRNA and treated with media without exosomes (No Exo), control exosomes (Co Exo) or 1X, 0.5X or 0.25X dose of RTT exosomes as described in Figure 5A. Quantitative analysis of apoptotic cells showed that there was no statistically significant increase in cell death with control or RTT exosome treatments compared to media control. B. Control exosomes rescue apoptotic effect of MeCP2 knockdown. Cultures were infected with lentivirus expressing shRNA for MeCP2 knockdown and treated with media (No Exo), 1X, 0.5X or 0.25X doses of control exosomes (Co Exo) or 10ng/ml IGF as described in Figure 4. MeCP2 knockdown (No Exo) increases apoptotic cells 3.59X ± 0.41 times compared to control (No Exo, shCO) exosome; p=0.0001). 1X control exosome treatment rescued apoptosis by reducing the number of apoptotic cells to 1.77X ± 0.33 (p=0.0001) compared to no exosome MECP2 knockdown, whereas other treatments had no effect. N=3 wells per group.

**Table S1: Quantitative proteomic analysis of control vs RTT exosomes.** Proteome of quantitative TMT analysis between control and RTT exosomes. 3611 proteins were identified in experiment 1 (tab 1) and 3019 proteins were identified in experiment 2 (tab 2). Average ratio of control/ RTT (column H, I) was calculated from three ratios (column D, E, F) from 6 replicates in each experiment.

**Table S2: Annotation of quantitative control vs RTT proteome for cellular location and protein type. Related to Figure 2, panel A.** Cellular location (column D) and protein type (column E) annotation using Ingenuity Pathway Analysis software for Proteomic dataset of 2572 proteins (tab 1) overlapping in experiments 1 and 2 and a subset of 237 proteins (tab 2) that were changed >1.5 fold between control and RTT exosomes.

**Table S3: PANTHER Overrepresentation Test of proteins changed >1.5 fold between control and RTT exosomes using the GO ‘biological processes’ complete annotation dataset. Related to Figure 2, panel B.** PANTHER Overrepresentation Test of the dataset of 237 proteins with >1.5 fold change using the GO ‘biological processes’ complete annotation dataset. The 237 candidates were classified into annotated GO biological process categories (column A) and compared with the normal human database (column B) to determine whether they are overrepresented or underrepresented (column E) and fold enrichment (column F) for a given GO biological process.

**Table S4: Canonical pathways from Ingenuity Pathway Analysis. Related to Figure 2, panel C and Figure 2S, panel C.** Canonical pathways from Ingenuity Pathway Analysis with -log p-value (column B) and overlap ratio (column C) for dataset of >1.5 fold changed (tab 1) and all (tab 2) differentially expressed proteins between control and Rett exosomes. The proteins from dataset that are represented in pathway are listed in column D.

**Table S5: Relative strength of downstream effects for biological functions and diseases in the nervous system. Related to Figure 2, panel D and Figure 2S, panel D.** Downstream effects for biological functions (column B and C) analysis by Ingenuity Pathway Analysis software in ‘Nervous System Development and Function’ category showing number of proteins from dataset (column D) and p-values (column E) for dataset of >1.5 fold changed (tab 1) and all (tab 2) differentially expressed proteins between control and Rett exosomes.

**Table S6: Comparison analysis of Canonical Pathways downstream effect for biological functions between experimental replicates. Related to Figure 2S, panel A and B.** Comparison analysis of canonical pathways using Ingenuity Pathway Analysis for experimental replicates, experiment 1 and 2 (tab 1) showing comparison of -log p-value (column C) and overlap ratio (column D) for each experimental replicate with proteins from dataset used for pathway listed in column E. Comparison of p-values (column B and C) between experimental replicates (tab 2) for downstream effects for biological functions analysis by Ingenuity Pathway Analysis software in ‘Nervous System Development and Function’ category.

## Star Methods

### CONTACT FOR REAGENT AND RESOURCE SHARING

Dr. Cline, 10550 North Torrey Pines Rd. La Jolla, CA 92037;

Dr.Muotri, 2880 Torrey Pines Scenic Drive, La Jolla, CA 92093. MC0695

### EXPERIMENTAL MODEL AND SUBJECT DETAILS

#### Human iPSC-derived neural cultures

To generate neural progenitor cells (NPCs), cells were differentiated and maintained as previously described (Chailangkarn et al., 2016; Marchetto et al., 2010). A male patient cell line with *MECP2* Q83X mutation, its male parental control and an isogenic rescue, corrected using CRISPR/Cas9 genome editing, were used in this study (Zhang et al., 2016). Two different clones of control (isogenic rescue or father of Q83X patient, WT83) iPSCs and RTT iPSCs were used in our studies to mitigate line-to-line variation. iPSCs lines maintained in mTESR media were switched to N2 (DMEM/F12 media supplemented with 1X N2 NeuroPlex Serum-Free Supplement (Gemini) with the dual SMAD inhibitors, 1 μM of dorsomorphin (Tocris) and 10 μM of SB431542 (Stemgent)) daily, for 48 hours. After two days, colonies were scraped off and cultured under agitation (95 rpm) as Embryoid Bodies (EB) for seven days using N2 media with dorsomorphin and SB431542. Media was changed every other day. EBs were then plated on Matrigel-coated dishes, and maintained in DMEM/F12 supplemented with 0.5X of N2 supplement, 0.5X Gem21 NeuroPlex Serum-Free Supplement (Gemini), 20ng/mL basic fibroblast growth factor (bFGF, LifeTechnologies) and 1% penicillin/streptomycin (P/S). After 7 days in culture, formed rosettes from the plated EBs were manually selected, gently dissociated with StemPro Accutase (LifeTechnologies) and plated onto 10 μg/mL poly-L-ornithine (Sigma)/ 5 μg/mL laminin Laminin (LifeTechnologies) coated plates. NPCs were maintained in DMEM/F12 with N2, Gem21, bFGF and P/S. NPCs were expanded as soon as confluent, using StemPro Accutase for 5 minutes at 37°C, centrifuged and replated with NGF with a 1:3 ratio in poly-L-ornithine /Laminin-coated plates. To induce cortical neuron differentiation, FGF was retrieved as previously described (Marchetto et al., 2010). The cortical neurons were allowed to differentiate for 4-8 weeks before exosome collection. The media was collected every 2 days for exosome isolation.

All cell lines used have been authenticated. All cell lines have tested negative for mycoplasma. Cytogenetic analysis was performed in all clones to evaluate correct cell karyotype (Children’s Hospital Los Angeles). In addition, all cell lines have been genotyped to ensure the presence of the mutation. The absence of the MeCP2 protein in the patient cell line has been verified by immunohistochemistry.

#### Human primary neural cultures

Human primary neurons were purchased from ScienCell, without identification of sex. Cells were thawed and plated according to the manufacturer’s instructions. After thawing, cells were counted and 20,000 cells per 96-well plate were plated in neuronal media onto 10 μg/mL poly-L-ornithine (Sigma)/ 5 μg/mL laminin (Life technologies)-coated plates. Cells were infected with lentivirus expressing shRNA targeted to *MECP2* (sh*MECP2*) or scrambled control shRNA (shControl) as described previously (Marchetto et al., 2010). For the rescue experiments, 10 ηg/mL of IGF1 was added to the cultures (Peprotech cat # 100-11R3).

#### Neurospheres for MEA

To generate functional neural networks, growing NPCs were dissociated with Accutase and plated on 6 well dishes (3-5 ×10^6^ cell per well) under shaker agitation (95 rpm) at 37°C. The media used was DMEM-F12 supplemented with 0.5X GEM21 (Gemini), 0.5X N2 (Gemini) and 20 ng/mL of bFGF (Life Technologies). The next day (day 0) neural fate was induced by removing bFGF and adding 10 μM Rock inhibitor Y-27632 (Tocris, USA). Two days later (day 2) media was changed to fresh media without Rock inhibitor. The differentiation took place for 2 weeks in suspension, with media change every 4 days. On day 15 of differentiation, these 3D neurospheres cultures were plated on poly-ornithine/laminin-pre-coated 12-well MEA plates (Axion) with DMEM-F12 media supplemented with 0.5X GEM21, 0.5X N2 and 1 %FBS. Media was changed every 3–4 days. After 1-2 weeks, media was replaced with Neurobasal supplemented with 1X B-27 and 1:400 Glutamax.

### METHOD DETAILS

Exosomes were enriched from media from hiPSC-derived neural cultures as described in Figure 1A (Thery et al., 2006). Fresh growth media was replaced in cultures 2 days prior to collecting exosomes. After 2 days, the culture media was harvested for exosome isolation and replaced with fresh growth media. For multiple treatments, exosomes were harvested from the media collected from same cultures every 2 days. The harvested media was centrifuged at 300g for 15 minutes at 4°C, the supernatant was collected and centrifuged at 2000g for 15 minutes at 4°C in a Beckman tabletop centrifuge. The second supernatant was further centrifuged using a Beckman ultracentrifuge at 10,000g for 45 min at 4°C. The supernatant was collected and centrifuged at 100,000g for 1 hour at 4°C to pellet exosomes. The exosome pellet was washed twice with D-PBS containing Ca^++^ and Mg^++^ (GIBCO) at 10,000g for 1 hour at 4°C followed by resuspension in normal growth media described above, at 1/500 of the original volume of media, resulting is 500X enrichment of exosomes. Exosomes were used for assays immediately after purification as we observed decreased bioactivity when exosomes were stored overnight at 4°C.

#### Proteomic Analysis

Twelve exosome preparations, half from RTT patient iPSC-derived neural cultures and half from isogenic control iPSC-derived neural cultures, were labeled with TMT isobaric tags for quantitative mass spectrometry (MS) analysis. Two independent MS experiments were performed each with 3 different RTT preparations and 3 different control preparations obtained from at least 3 independent batches of neuronal differentiations of NPCs for each line, and thus, 6 biological replicates were analyzed together in each MS experiment. We detected 3611 and 3019 annotated proteins in experiment 1 and 2, respectively, with 2572 proteins (62% overlap) detected in both MS experiments. Of these, 873 and 789 proteins in experiments 1 and 2, respectively, were annotated for function in neurons with 739 detected in both (80% overlap). We compared these datasets to identify ‘cellular pathways’ and ‘downstream effects on biological functions in the nervous system’ using the manually curated Ingenuity database (Calvano et al., 2005). The two experiments provided very similar prediction results regarding ‘cellular pathways’ with *p*-values and overlap ratios (the fraction of proteins annotated for each pathway that were detected in the experimental dataset) ranging from 10^−56^ to 10^−21^ for *p*-values and 0.65 to 0.44 for overlap ratios for the top 10 pathways (Figure S2A). Similarly, analysis of ‘downstream effects on biological function in nervous system’ demonstrated excellent reproducibility between the two experiments with p-values ranging from 10^−73^ to 10^−37^ for the top 10 categories (Figure S2B). The analysis performed on the 2572 overlapping proteins was largely similar to results from individual experiments (Figure S2C-D).

The TMT labeling was performed according to the manufacturer’s instructions (Thermo Fisher Scientific, Rockford, IL) with modifications as previously described (Rauniyar et al., 2013). Prior to loading on a MudPIT trapping column, the labeled peptide mixture was diluted five times with buffer A (5% acetonitrile/0.1% formic acid) and then centrifuged at 12,000 x g for 30 min to remove particulates.

LC-MS/MS analysis was performed on an LTQ Orbitrap Velos instrument (Thermo Scientific, San Jose, CA, USA) interfaced at the front end with a quaternary HP 1100 series HPLC pump (Agilent Technology, Santa Clara, CA, USA) using an in-house built electrospray stage. The SEQUEST search and DTASelect analysis were performed identically as previously described (Rauniyar et al., 2013). To confirm the TMT labeling efficiency, the spectra were also searched without these modifications. Census, a software tool for quantitative proteomic analysis (Park et al., 2008), was used to extract the relative intensities of reporter ions for each peptide from the MS2 spectra. Analysis of TMT quantification datasets with Census was performed as previously described (Rauniyar et al., 2013).

Ingenuity Pathway Analysis software version 01-08 (QIAGEN Bioinformatics) was used for annotations and pathway analyses. PANTHER Overrepresentation Test (release 20170413) was performed using GO Ontology database (released 2017-04-24) using Homo Sapiens database as a reference list and GO biological processes complete as an annotation dataset. Bonferroni correction for multiple testing was applied for statistics.

#### Assays

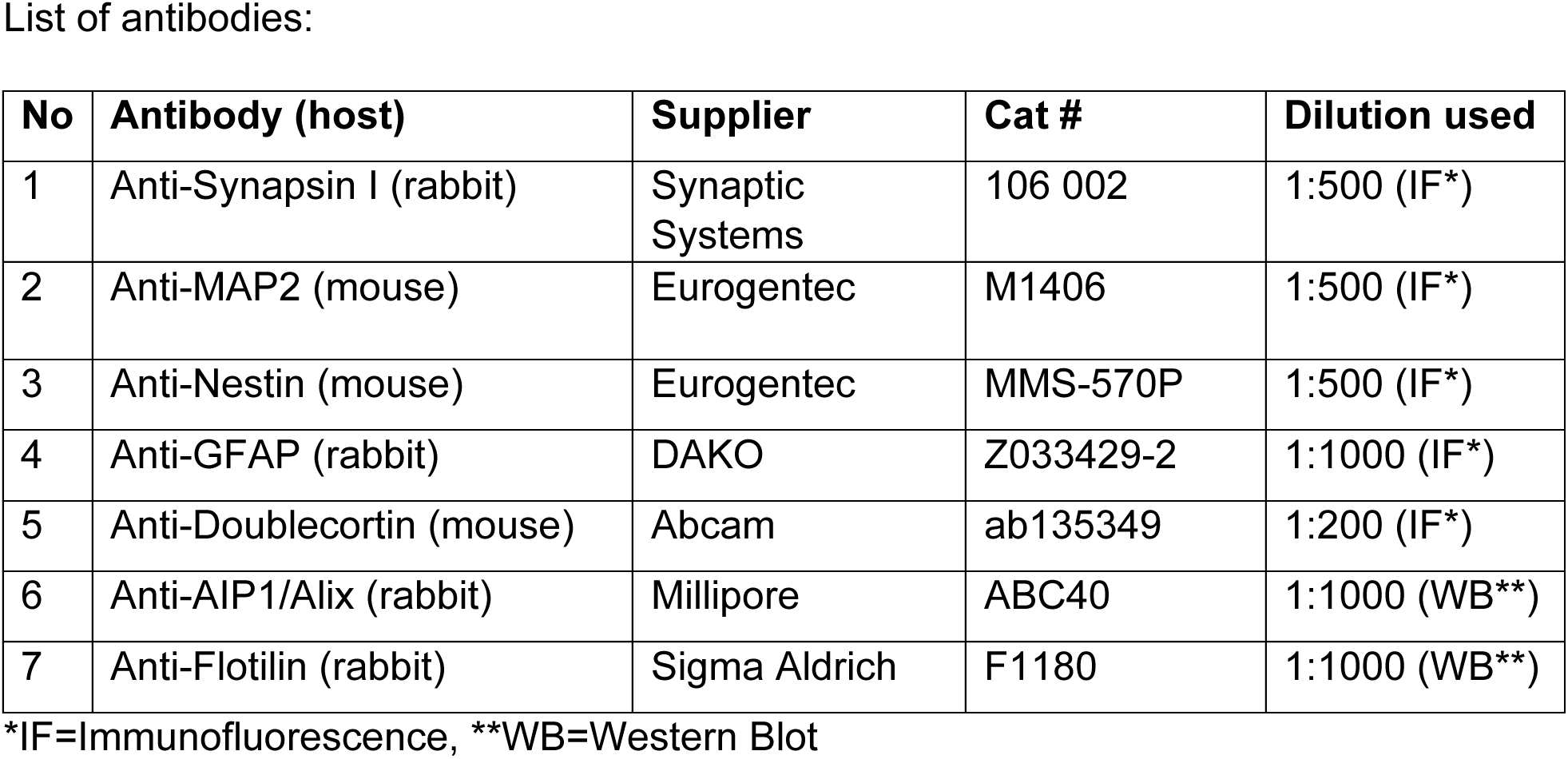

##### Exosome/EdU treatments

Purified control or RTT exosomes were resuspended in fresh growth media and added to lentivirus infected human primary neural cultures, iPSC-derived neural cultures by completely replacing old media. For neurospheres the exosomes were resuspended as 10X concentrated stock of doses and added onto neutrosphere cultures without replacing media to achieve desired dose concentration. Exosome treatment was performed every second day, 2-4 times, as described in the schematics in the figures. For EdU labeling experiments, human primary neural cultures were treated with control or RTT exosomes on DIV 5 and 7. On day 7 a 2-hour pulse of 10 μM EdU (ThermoFisher Scientific) was provided prior to addition of exosomes.

##### Immunolabeling, microscopy and image analysis

After treatments, cultures were fixed in 4% paraformaldehyde at room temperature (RT) for 15 min and then permeabilized with 0.5% Triton X-100 in D-PBS with Ca^++^ and Mg^++^ at RT for 5 min. Cells were then blocked in 5% normal goat serum in D-PBS with Ca^2+^ and Mg^2+^ at RT for 1 hr before incubation with primary antibody (list of primary antibodies provided in supplemental data) diluted in 5% normal goat serum overnight at 4°C. After three washes with D-PBS containing Ca^++^ and Mg^++^, cells were incubated with Alexa 488, Alexa 568 or Alexa 633 Goat anti-Mouse IgG, Goat anti-Rabbit IgG or Goat anti-Chicken IgY (depending on host for primary antibodies as listed in supplemental data), (H+L) highly cross-adsorbed secondary antibodies (Invitrogen/ThermoFisher) followed by 3 washes with D-PBS containing Ca^++^ and Mg^++^. During the last wash, the cultures were labeled with DAPI solution (3 ng/mL) for 10 minutes and mounted in Prolong Gold mounting media (Invitrogen/ThermoFisher). Each experimental condition was blinded before imaging. Imaging was performed with Plan Apo 20X objective with 0.75 NA on a Zeiss LSM 710 or Nikon A1 confocal system with Andor iXon EMCCD camera system. Images were collected from randomized areas in each dish and analyzed using Metamorph software (Molecular Devices). Synaptic puncta analysis was performed using a custom Image J plugin. A trimming procedure was applied to isolate and segregate individual puncta based on the inclusion of pixels with intensities greater than 0.3 of maximal intensity within a unit of connected pixels having intensity greater than the set threshold. This was achieved by an iterative procedure using a step size (0.01) for each iteration, which limits for the number of iterations of the trimming procedure. The resultant image consists of a background subtracted image containing a set of well-defined puncta and provides quantification of net intensity per spot and total pixel area per spot. Apoptosis was quantified using DAPI nuclear labeling. Cells with condensed and blebbing or fragmented nuclei were scored as apoptotic. The fraction of apoptotic cells was calculated from the count of total DAPI-labeled nuclei.

##### Multielectrode Array Recordings

We recorded spontaneous action potential activity using the Axion Biosystems. RTT or control exosomes were resuspended in neurosphere media and were added directly on top of RTT or control neurospheres that were plated on the MEAs. Neurosphere activity was recorded before and 24 hours after each exosome addition. Exosomes were added every 2nd day to neurospheres for 1-2 weeks. Only MEA channels with similar number and density of cells plated, and with more than 10 spikes in a 5 min interval were used in the analysis. Recordings were analyzed with the software Neuroexplorer (Nex Technologies, Madison, AL, USA) and MATLAB-based Neural Metric Tool Software Axis (Axion Biosystems). Raster plots were generated using Neuroexplorer.

### QUANTIFICATION AND STATISTICAL ANALYSIS

Statistics was performed using Microsoft Excel, Statistics Online Computational Resource (SOCR) of UCLA (http://www.socr.ucla.edu/SOCR.html) and GraphPad Prism 7.03. For statistics 2 way ANOVA with Bonferroni correction applied for multiple comparisons or independent sample Wilcoxon rank sum test for pairwise comparisons was performed. P values are provided in figure legends. For cumulative frequency distribution, the Kolmogorov-Smirnoff test was performed. D-stat, the absolute maximum difference between two cumulative frequency distribution function, and p-values are provided in figure legend.

